# Multiplexed mRNA analysis of brain-derived extracellular vesicles upon experimental stroke in mice reveals increased mRNA content related to inflammation and recovery processes

**DOI:** 10.1101/2021.12.09.471913

**Authors:** Annika Bub, Santra Brenna, Malik Alawi, Paul Kügler, Yuqi Gui, Oliver Kretz, Hermann Altmeppen, Tim Magnus, Berta Puig

**Author notes:** shared first authorship (Annika Bub, Santra Brenna).

## Abstract

Extracellular vesicles (EVs) are lipid bilayer enclosed structures that not only represent a newly discovered means for cell-to-cell communication but may also serve as promising disease biomarkers and therapeutic tools. Apart from proteins, lipids, and metabolites, EVs can deliver genetic information such as mRNA eliciting a response in the recipient cells. In the present study, we have analyzed the mRNA content of brain-derived EVs (BDEVs) isolated 72 hours after experimental stroke in mice and compared them to controls (shams) using the nCounter® Nanostring panels, with or without prior RNA isolation from BDEVs. We found that both panels show similar results when comparing upregulated mRNA in stroke. Notably, the higher upregulated mRNAs were related to processes of stress and immune system responses, but also to anatomical structure development, cell differentiation, and extracellular matrix organization, indicating that regenerative mechanisms are already taking place at this time-point. The five top overexpressed mRNAs in stroke mice compared to shams were confirmed by RT-qPCR and, interestingly, were found to be present as full-length open-reading frame in BDEVs. We could reveal that the majority of the mRNA cargo in BDEVs was of microglial origin and probably predominantly present in small BDEVs (≤ 200 nm in diameter). However, the EV population with the highest increase in the total BDEVs pool at 72 h after stroke was of oligodendrocytic origin. Our study shows that nCounter® panels are a good tool to study mRNA content in tissue-derived EVs as they can be carried out even without previous mRNA isolation and that the mRNA cargo of BDEVs indicates their participation in inflammatory but also recovery processes after stroke.

## INTRODUCTION

Extracellular vesicles (EVs) are lipid bilayer particles secreted by all types of cells that play an important role in communication among cells as they can transfer proteins, lipids, and genetic material to recipient cells, even over long distances^1,2^. Among the genetic information contained in EVs, different types of RNA cargoes have been identified such as non-coding RNAs (miRNA, tRNA, siRNA, YRNA, lncRNA, circRNA), and fragmented as well as intact mRNAs^3,4^. The loading of RNAs into EVs is not random as earlier studies support selective incorporation, reflecting the type and the physiological state of the parent cells^5,6^. Moreover, several studies have reported that different types of EVs originating from the same cell type contain differentially sorted miRNAs and mRNAs^7-9^.

The mRNAs in EVs are transferred to and translated into protein in recipient cells^10-14^. Other types of RNAs such as fragmented non-coding RNAs and miRNAs have been shown to play an important role in cancer development^14-18^. However, many of the physiological consequences of mRNA and non-coding RNAs contained in EVs once taken up by recipient cells are still unclear.

Ischemic stroke is the world-leading cause of sustained disability. It has a complex pathophysiology involving the reaction of both, brain and infiltrating immune cells, the latter due to the breakdown of the blood-brain barrier (BBB)^19^. After the blockage of an artery, a transient lack of glucose and oxygen at the core of the stroke triggers necrotic cell death in neurons which release their content to the extracellular space, generating Danger Associated Molecular Patterns (DAMPs, such as ATP). DAMPs activate microglia and astrocytes, triggering an immune response. Surrounding the ischemic core, there is a hypoperfused area, the penumbra, where cells are still metabolically active and could potentially be rescued within a critical timeframe^20,21^, probably implying a better outcome for the patient^22,23^.

Several studies support the idea that EVs play an important role in deciding the neuronal fate under stress conditions and probably the same applies to the neuronal outcome in the penumbra after stroke. On the one hand, *in vitro* studies showed that extracellular ATP stimulates microglia to release EVs containing an altered proteome compared to steady-state conditions, triggering an inflammatory phenotype in astrocytes^24^. ATP released by astrocytes also stimulates the release of microglial EVs containing the proinflammatory cytokine IL-1β^25,26^. On the other hand, EVs derived from non-stimulated astrocytes can rescue neurons under ischemic conditions, a function that depends on the cellular prion protein (PrP^C^)^27^. Furthermore, EVs derived from oligodendrocytes increase the viability of neurons subjected to nutrient deprivation^28-30^. Interestingly, a recent study proposed that neurons release “help-me” signals through EVs^31^. The authors showed that EVs containing miRNA-98 released by neurons were taken up by microglia *in vitro* and *in vivo* and, consequently, microglial phagocytosis of stressed but still salvageable neurons was decreased in the penumbra after stroke. Thus, to study the intercellular communication after stroke through EVs is of utmost importance to (i) understand the underlying pathophysiological mechanisms, (ii) identify novel biomarkers, and (iii) develop new therapeutic approaches.

Our present study aimed to investigate if and how the mRNA content in BDEVs changes after stroke compared to sham. To this end, we applied a targeted approach using the nCounter® Neuropathology Panel allowing for the simultaneous assessment of 770 genes related to diverse aspects of neurodegeneration such as neurotransmission, neuron-glia interaction, and neuroinflammation, among others. As these panels can also be used without previous mRNA extraction, another aim was to investigate if this was also applicable to the study of tissue-derived EVs, which would represent a technical advantage as it would eliminate steps in the protocol, thus reducing sample loss and decreasing variability. We subjected mice to transient Middle Cerebral Artery Occlusion (tMCAO), a widely stablished mouse model of stroke, and explored the mRNA content of BDEVs at 72 h after reperfusion, when recovery processes may start to take place. We show that (i) the nCounter® panels can be used for EVs analysis bypassing the mRNA isolation, obtaining similar results to the panels incubated with previously isolated mRNA; (ii) the highest increase in BDEVs from tMCAO mice was observed for mRNAs related to inflammatory and recovery processes with (iii) mRNA top hits being present as full-length and mostly contained in small EVs (≤200nm); (iv) the majority of highly upregulated mRNAs in BDEVs are released by microglia at this time point even though (v), the most upregulated contributors to the whole BDEV pool at 72 h after stroke are oligodendrocytes. To the best of our knowledge, this is the first report of mRNA analysis from BDEVs. From a technical point, the present study shows that nCounter® panels are a convenient and reliable method to study EVs as they can perform without prior mRNA isolation. We also show that full-length mRNAs with possible implications in inflammatory and regenerative processes are increasingly shuttled in EVs after stroke, revealing conceivable regulatory roles in stroke pathophysiology.

## MATERIAL AND METHODS

### Ethics statement

All animal experiments were approved by the local animal care committee (*Behörde für Gesundheit und Verbraucherschutz, Veterinärwesen und Lebensmittelsicherheit of the Freie und Hansestadt Hamburg*, project number N045/2018), and performed following the guidelines of the animal facility of the University Medical Center Hamburg-Eppendorf.

Mice used for this study were kept under a 12h dark-light cycle with *ad libitum* access to food and water.

### Transient middle cerebral artery occlusion (tMCAO) procedure

12-18 weeks old male C57BL/6 mice were used for the experiments. Mice were anesthetized and tMCAO was performed as described previously^19^. Briefly, the temporary occlusion of the middle cerebral artery was achieved by inserting a 6.0 nylon filament (602312PK10, Doccol) for 40 min. Control (sham) animals were also anesthetized, and their arteries were exposed but not occluded. Mice were euthanized 72 h after the tMCAO procedure. Ipsilateral and contralateral hemispheres were stored separately at –80 °C. For the present study, only the ipsilateral hemispheres were used.

### EVs isolation

EVs were isolated from the ipsilateral hemisphere of mice from the stroke or sham group as described previously^32^. Briefly, frozen brains were slightly thawed in Hibernate E media (Gibco), finely chopped, and digested with 75U/mL of Collagenase type III (Worthington) in Hibernate E (800μL/ 100mg tissue) for 20 min in a shaking water bath at 37°C. Digestion was stopped using protease inhibitors (Roche). Samples were centrifuged at 300×*g* for 5 min at 4°C, the supernatant was collected and further centrifuged at 2,000×*g* for 10 min at 4°C. The resultant supernatant was further centrifuged at 10.000×*g* for 30 min at 4°C. For some experiments (“filtered samples”, F), the resultant supernatant was passed through a 0.22µm cellulose-acetate filter (Whatman) or directly used in the next step (“non-filtered samples”, NF). The supernatant was layered on top of a sucrose gradient (2.5M, 1.3M, 0.6M) and centrifuged at 180,000 ×*g* (corresponding to 31,800 rpm in a SW40Ti rotor) for 3h at 4°C. Six fractions were collected, diluted in PBS (Gibco), and further centrifuged at 100,000×*g* (24,000 rpm in SW40Ti rotor) for 1h at 4°C. After the initial characterization, fractions 2,3, and 4 were pooled before dilution in PBS. The pellet was stored at -20°C for further RNA isolation or resuspended in RIPA buffer (50 mM Tris-HCl pH=7.4, 150 mM NaCl, 1% NP40, 0.5% Na-Deoxycholate and 0.1% SDS) or PBS with protease and phosphatase inhibitors (Roche) to be used for western blot or NTA analysis, respectively.

### SDS-PAGE and western blot

EVs samples in RIPA were mixed with 4× NuPage LDS Sample buffer (Invitrogen) and 10× NuPage Sample reducing agent (Invitrogen) and heated at 70°C for 10 min. For EVs characterization, the same volume for the 6 fractions was loaded (15 µL) in NuPage 10% Bis-Tris precast gels and run at 150V. Proteins were then transferred onto nitrocellulose membrane (LI-COR) at 400 mA for 1h. Total protein content was visualized by staining the membranes with Revert Total Protein Stain Kit (LI-COR), as described by the manufacturer’s instructions. To block unspecific binding, membranes were incubated with 1× RotiBlock (Roth) in TBS for 1h at room temperature while shaking and incubated overnight at 4°C with the following primary antibodies: ADAM10 (1:1,000; EPR5622), Alix (1:500; #2171;CST), CD81 (1:1,000; #10037; Cell Signaling), Flotilin-1 (1:1,000; #610820; BD Biosciences), or GM130 (1:1,000; #61082; BD Biosciences). After incubation, membranes were washed with TBST, incubated for 1h with the corresponding secondary antibody (1:1,000 Anti-mouse IgG,#7076/Anti-rabbit IgG, #7074; both from Cell Signaling), washed again with TBST, and developed with Super Signal West Femto Substrate (ThermoFisher). The chemiluminescence reaction was detected with a ChemiDoc Imaging Station (BioRad).

For cellular markers characterization, 3 µg of proteins measured with the Micro BCA Protein Assay Kit (Thermo Scientific) following the instructions of the supplier, were mixed with 4× loading buffer (250mM Tris-HCl, 8% SDS, 40% glycerol, 20% β-mercaptoethanol, 0.008% Bromophenol Blue, pH 6.8), boiled for 5 min at 95°C, loaded onto a 10% Bis/Tris acrylamide gel and subjected to electrophoresis as described above. The following antibodies were used: CD40 (1:500, NB100-56127SS; Novusbio), CNP(1:1,000, C5922; Sigma), EAAT1 (1:500, NB100-1869SS; Novusbio), EAAT2 (1:500, NBP1-20136SS; Novusbio), NCAM (1:1,000, #99746; Cell Signaling), P2Y12 (1:500, #11976-1-AP; Proteintech), PLP (1:1,000, NB100-74503; Novusbio), Synapsin-1 (1:1,000, #106103; Synaptic Systems). After overnight incubation while shaking at 4°C, the detection was performed as described above.

### RNA isolation

RNA was isolated from the EVs using the Qiagen RNeasy Plus Micro Kit (Qiagen). Briefly, EV pellets from fractions 2, 3, and 4 were resuspended each in 117 µL of Lysis Buffer RLT Plus (Qiagen), vortexed, and pooled together (total of 350 µL as recommended by the supplier) and RNA was isolated following the manufacturer’s instructions. The quality and purity of the isolated RNA were checked using the Agilent 2100 Bioanalyzer following the instructions of the supplier.

### Gene expression analysis with nCounter® panels

Gene expression analysis was performed by using the NanoString nCounter® Neuropathology panel (#XT-CSO-MNROP1-12, NanoString Technologies). Two panels were used: one loaded with RNA isolated from BDEVs (that were filtered during the preparation of EVs); and another one where all the samples bypassed the RNA isolation (“non-isolated”, NI) but some were filtered during the BDEVs preparation (F) and the others were not filtered (NF) during preparation.

To run the first panel, 50 ng of previously isolated RNA measured by Qubit™ RNA High Sensitivity Assay Kit (Thermo Fisher) using the 3.0 QuBit Fluorometer, were mixed with RNAse–free water (Qiagen) up to 5 µL. Samples were hybridized for 18 h and mixed with 15 µL of RNAse-free water to be loaded on the nCounter® Sprint Cartridge (#LBL-10038-01, NanoStringTechnologies), following the instructions of the company. The analysis was run for 6 h.

For direct loading of EV lysates (i.e., no previous RNA isolation, NI), frozen EVs were resuspended in RLT lysis buffer and RNAse-free water in a ratio of 1:3 and loaded based on protein concentration measured by Micro BCA Protein Assay Kit (Thermo Scientific). 2,8 μg of proteins were loaded.

### Reverse Transcription and quantitative PCR (RT-qPCR)

The cDNA was synthesized from the BDEVs’ isolated RNA using Maxima First Strand cDNA Synthesis Kit for RT-qPCR (Thermo Scientific), following the supplier’s instructions. The resulting cDNA was loaded in a 1:15 dilution with a master mix (Probe qPCR MM No ROX; Thermo Scientific) with the following TaqMan Probes: Hmox1 (#Mm00516005_m1), Fcrls (#Mm00472833_m1), Cd44 (#Mm01277161_m1), C1qb (#Mm01179619_m1), Gfap (#Mm01253033_m1) all from Thermo Scientific. Asb7 (#Mm01318985_m1) and Fam104a (#Mm01245127_g1), both from Thermo Scientific, were used as housekeeping genes based on the nCounter® analysis. The reaction was performed using LightCycler 96 (Roche) with the following conditions: 50°C for 120 s, 95°C for 600 s, followed by 45 cycles of 95°C for 10 s, and 60°C for 30 s and further cooling of 37°C for 30 s. Samples were run in triplicates. Differential expression was analyzed using the 2^-ΔΔCT^ method. ΔCT was calculated by subtracting the arithmetic means of the CT values of our two housekeeping genes from the CT values of the mRNA of interest. ΔΔCT was calculated by subtracting the average ΔCT of the sham samples from the ΔCT of the stroke samples. The fold change (FC) was calculated as 2^-ΔΔCT.^

### PCR

cDNA was synthesized as above and 5µL of the resulting EVs cDNA (concentration not measured as the PCR was only intended to be qualitative) or 1 µL of total WT brain cDNA (positive control) or 1 µL of water (negative control) were mixed with 1× Master Mix: dNTPs 0.4 mM, primers 0.4 µM, Dream Taq polymerase 1U, 1× Dream Taq Green Buffer (all from Thermo Fischer) and water. The following primers were used:

<u>Hmox1</u>: 5’
sATGGAGCGTCCACAGCC 3’(sense), 3’ GGCATAAATTCCCACTGCCAC 5’(antisense) <u>C1qa</u>: 5’ATGGAGACCTCTCAGGGATGG 3’(sense), 3’TCAGGCCGAGGGGAAAATGA 5’(antisense); <u>C1qb</u>:5’TGAAGACACAGTGGGGTGAGG 3’(sense), 3’TACGCATCCATGTCAGGGAAAA 5’(antisense); <u>C1qc</u>: 5’ATGGTCGTTGGACCCAGTTG 3’(sense), 3’CTAGTCGGGAAACAGTAGGAAAC 5’(antisense); <u>Gfap</u>: 5’ATGGAGCGGAGACGCATCA 3’(sense), 3’ACATCACCACGTCCTTGTGC 5’(antisense); and <u>Cd44</u>: 5’GTTTTGGTGGCACACAGCTT 3’(sense), 3’CAGATTCCGGGTCTCGTCAG 5’(antisense).

The PCR reaction was performed as follows: 95°C for 5 min; 95°C for 45 s and 61°C for 45 s, and 72°C for 1 min (×35 cycles); 72°C for 5 min. The samples were loaded in a 1.5% Agarose gel mixed with Roti-GelStain (Roth) and run at 120 V for 40 min. The gel was then stained with ethidium bromide (Fluka) and imaged with UVP UVsolo touch (Analytik Jena).

### Nanoparticle Tracking Analysis (NTA)

Pellets resulting from the EVs isolation were resuspended in 30 µL PBS and 1 µL of this suspension was diluted in PBS at 1:500, and the resulting 500 µL were then loaded into the sample chamber of the LM10 unit (Nanosight, Amesbury, UK). Samples were recorded with ten videos, each 10 s long. Data analysis was performed by NTA 3.0 software (Nanosight) with the following software settings: detection threshold = 6, screen gain = 2.

### Transmission Electron Microscopy

For TEM, extracellular vesicle pellets were re-suspended in 0.1M PBS. Drops of these suspensions were placed on parafilm. Carbon-coated copper meshed grids (Plano, Germany) were placed on the drops for 5 minutes for probe adsorption. After five minutes of fixation on drops of 1% glutaraldehyde (Roth, Germany) grids were washed 4 times for 30 seconds and negative contrasted using 1% uranyl acetate. Grids were air-dried and analyzed using a Philips CM 100 transmission electron microscope.

### Analysis of the nCounter® panel data, GO terms enrichment, and cell types of origin

For each of the three datasets, genes with raw expression scores not exceeding the average plus two standard deviations of all corresponding negative control probes, in at least one sham and one stroke sample, were removed from the analysis. Normalization was based on the ten housekeeping mRNAs present on the panel. Differential expression analysis was carried out with DESeq2^33^. A mRNA was considered differentially expressed if the corresponding absolute log2-fold change (log2FC) was larger or equal to 1 and the False Discovery Rate (FDR) was smaller or equal to 0.1. To perform overrepresentation analysis, clusterProfiler^34^ was used in combination with GOslim terms^35^.

A gene expression matrix for the data set ‘Whole Cortex & Hippocampus - 10X Genomics (2020) was obtained from the Allen Mouse Brain Atlas^36^. To generate this matrix we used the mRNA values corresponding to log2FCs greater or equal to 2 and FDRs smaller or equal than 0.1 in the panel with samples I+F.

### Statistical analysis

To analyze the differences between sham and strokes in the western blot and for the NTA we used GraphPad Prism 8, applying non-parametric Mann-Whitney U test in both cases. Statistical significance was considered for \**p* < 0.05, \*\**p* < 0.005, and \*\*\**p* < 0.001. Values are given as a mean± standard error of the mean (SEM). The exact *p*-values are given in the main text.

## RESULTS

### Characterization of BDEVs and mRNA isolation

An overview of our experimental strategy and workflow concerning the respective data in the main figures is shown in Fig. 1A. EVs from mouse brains were isolated as described before^32^ and characterized by western blot (WB), nanoparticle tracking analysis (NTA), and electron microscopy (EM) following the MISEV 2018 guidelines^37^. As shown in Fig. 1B, six fractions were obtained after sucrose gradient centrifugation. Fractions 2, and 3 were enriched in common EV markers such as flotillin, CD81, mature ADAM10, 14-3-3, and Alix (with the latter two being cytosolic proteins, indicating EV integrity). GM130, a membrane protein of the cis-Golgi apparatus was used here as a marker of contamination with intracellular organelles and was not present in any of the six fractions. For subsequent experiments, we pooled fractions 2, 3, and 4 (as in the latter fraction flotillin and CD81 were also present) as the “EVs fraction”. NTA measurements of this EV fraction revealed that the number of EVs was not significantly elevated in the tMCAO brains compared to shams, although there was a clear tendency towards an increase (mean value shams: 5.5×10^11^ ± 1.77×10^11^; mean value tMCAOs: 1.07×10^12^ ± .,82×10^11^; Fig. 1C; n=6 for each group). EM pictures showed that EVs of different sizes were enriched in the pooled pellets (Fig. 1D, black arrowheads). Some structures with a more squared profile were also identified (white asterisks) probably indicating some degree of contamination, yet no major differences were observed between EVs isolated from tMCAO brains compared to shams). We then proceeded to isolate RNA from the EVs fraction and analyzed it with Bioanalyzer. Most of the RNA contained in the EVs from shams or stroke brains corresponded to small RNAs of less than 1,000 nucleotides (nt), although some bands appeared between 1,000 and 4,000 nt as well. Very low amounts of the ribosomal RNA (rRNA) subunit 18S and no detectable levels of the 28S rRNA subunit were observed (Fig. 1E). A representative RNA electropherogram is shown in Fig. 1F confirming that the majority of RNA is less than 1,000 nt with only a few ranging between 1,000 and 4,000 nt. The mean RNA concentration for shams was 23.06 ± 2.053 ng/µL) whereas for strokes this was 22.06 ± 3.64 ng/µL (n=3 per condition) as measured with QuBit™ RNA High Sensitivity Assay.

**Figure 1.**
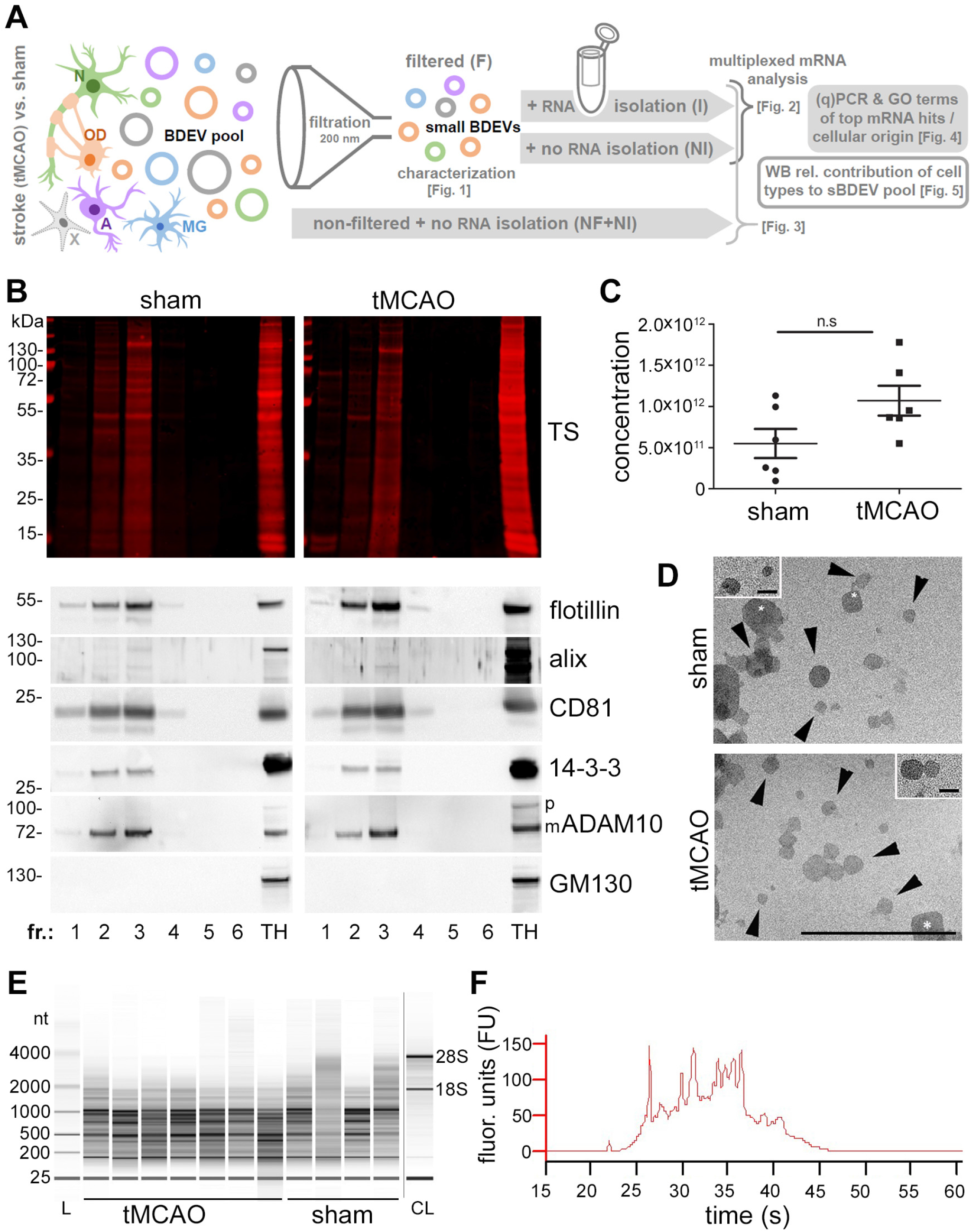
BDEVs characterization and mRNA isolation. (A) Overview of the research strategy and experimental workflow followed in the present study. Different cell types, including neurons (N, green), oligodendrocytes (OD, orange), astrocytes (A, pink), microglia (MG, blue), and others (X, grey) contribute to the EV pool in brain. EVs were purified from the brains of mice 72 hours after experimental stroke (tMCAO) or a control procedure (sham). Small brain-derived EVs (sBDEVs) were obtained upon filtration (F) and characterized. The mRNA content of sBDEVs was assessed in detail for stroke and sham samples and compared with (I) or without (NI) a previous RNA isolation step. Moreover, a comparison was performed between filtered (F, sBDEVs) and non-filtered (NF) BDEVs (with the latter population also containing larger EV species). Lastly, changes in the relative contribution of different cell types to the EV pool upon stroke were assessed. (B) Western blot characterization of the six fractions obtained after sucrose gradient centrifugation. Fractions 2 and 3 are labeled with antibodies against EV markers flotillin, CD81, and 14-3-3 indicating enrichment of EVs. Moreover, presence in the same fractions of Alix and mature (m) ADAM10 indicates enrichment in exosomes. CD81 and flotillin are also found in fraction 4, therefore fractions 2, 3, and 4 were pooled for the subsequent experiments. The Golgi protein GM130 is absent in the BDEVs fractions indicating a lack of contamination with intracellular organelles. TS is total protein staining. TH is a total brain homogenate used for comparison purposes. (C) Nanoparticle tracking analysis (NTA) of pooled BDEVs fractions (n=6). Values are given in the main text. (D) Electron microscopy of BDEVs. Arrowheads point towards BDEVs, whereas the white asterisks mark structures that, for the shape, are not assignable to EVs and most likely represent some minor contamination by cell membrane fragments. The scale bar is 500 nm, 100 nm on the insert. (E) Example of the BDEVs-RNA profile obtained with the Bioanalyzer. Most of the RNA is under 1,000 nt in both, tMCAO and sham BDEVs. CL is a cell lysate, used for comparison purposes as it shows the two main rRNAs (18S and 28S), which are mostly absent in BDEVs. (F) Representative electropherogram obtained with the Bioanalyzer showing the fluorescent units (FU) on the Y-axis and the migration time (in seconds, s) on the X-axis.

### Similar BDEVs’ mRNA profiles are obtained with the nCounter® panels with and without previous mRNA extraction, and with and without a filtration step during preparation of BDEVs

Isolated mRNA from EVs purified from the ipsilateral hemisphere of either sham (n=3) or tMCAO (n=3) mice were hybridized with the nCounter® Neuropathology panel from Nanostring which allows for multiplexed detection and quantification of 770 genes related to several aspects of neurodegeneration. Of interest for the studies of EVs’ mRNA, this method can detect low abundant mRNA^38^. To analyze the data, we used two criteria, a stringent one, focusing on those mRNAs that showed an absolute log2 fold-change (log2FC) larger or equal to 2 (i.e., increase/decrease), and a broader criteria also considering mRNAs increased/decreased with an absolute log2FC larger or equal to 1, respectively. The false discovery rate (FDR) in each case was required to be below or equal to 0.1. Out of the 770 genes present in the panel, 94 genes were significantly upregulated (log2FC≥1) and 3 were significantly downregulated (log2FC≤-1) in the tMCAO BDEVs compared to shams (Fig. 2A; Suppl. Table 1). A log2FC≥2 cut-off resulted in 31 mRNAs showing upregulation with Hmox1 being the highest hit with a log2FC of 3.73 (i.e., about 13-fold upregulated). However, the mRNA with the highest counts significantly upregulated in tMCAO (indicating a strong presence in BDEVs), was for Gfap (Fig.2B, Suppl. Table 1). No downregulated mRNA presented a log2FC≤-2. Principal component analysis (PCA) showed that samples of either sham or tMCAO cluster together, respectively, indicating differential mRNA expression between the two experimental groups (Suppl. Fig. 1).

**Figure 2.**
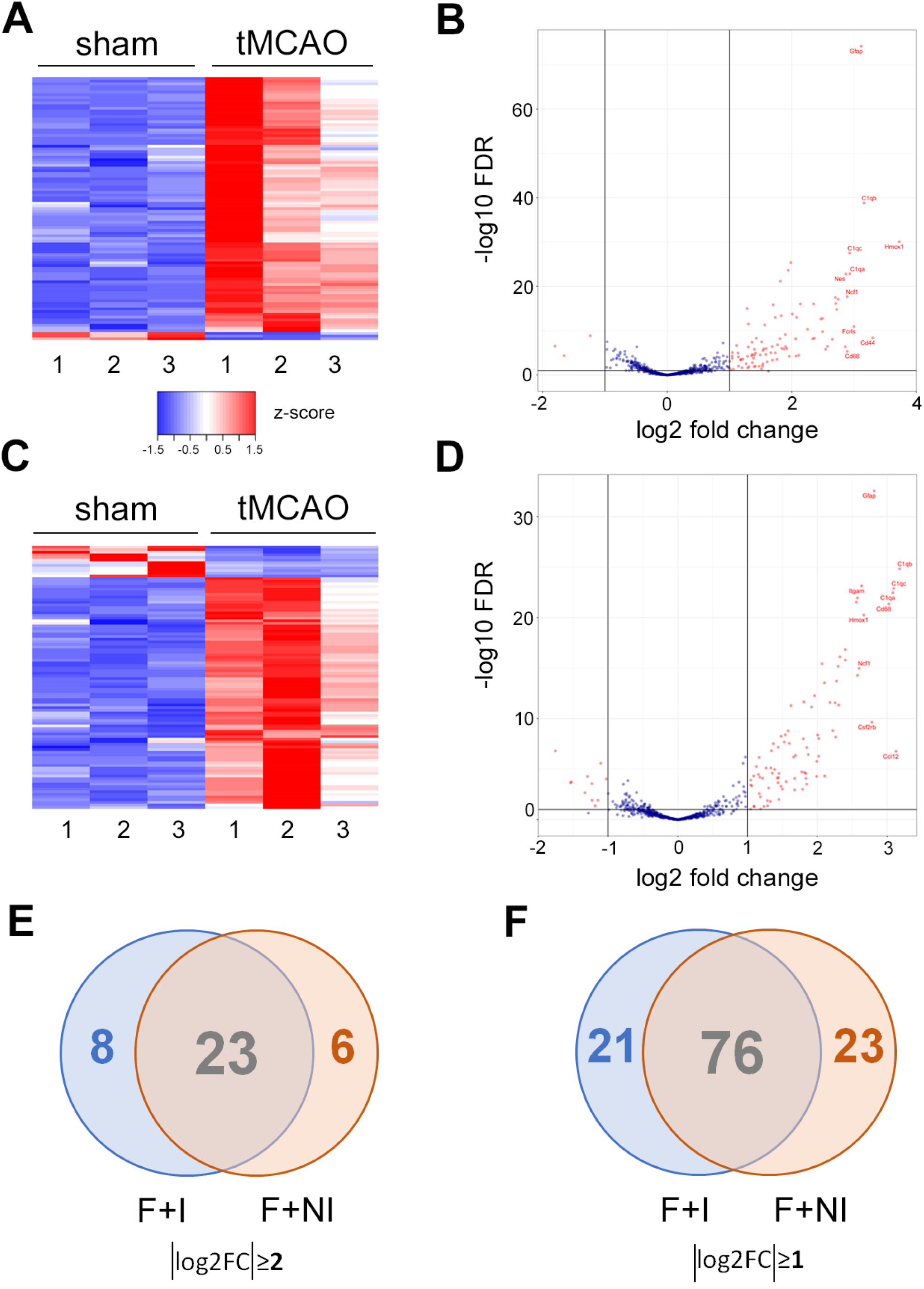
No major differences for the highest upregulated mRNAs in tMCAO sBDEVs compared to shams are observed with and without mRNA isolation. (A) Heat map showing the significantly up- (log2FC≥1) and downregulated (log2FC≤-1) mRNAs found in tMCAO mice compared to shams (n=3 mice per group) using the nCounter® Neuropathology panel with previous mRNA isolation (I) from sBDEVs. (B) Volcano plot for the same results as in (A) displaying the names of the ten significantly differentially expressed mRNAs with the highest log2FC in sBDEVs in tMCAO mice compared to shams. On the X-axis the log2FC is plotted while in the Y-axis shows the -log10 FDR. (C) Heat map showing the significantly up- and downregulated (absolut log2FC≥ 1) mRNAs in BDEVs from tMCAO compared to shams (n=3 per group) using the nCounter® panel without mRNA isolation. (D) Volcano plot for the same mRNAs as in (C) displaying the names of the ten significantly differentially expressed mRNAs in sBDEVs in tMCAO mice compared to shams. The X-axis shows the log2FC, the Y-axis the -log10 FDR. (E) Overview of the most up- and downregulated mRNAs (absolut log2FC≥2) with both types of preparations (i.e., with mRNA isolation (I) and without mRNA isolation (NI)) displayed in a Venn diagram showing the number of the commonly shared and unique mRNAs for each panel. (F) Venn diagram for the different sample conditions as in (E) but with a cut-off of absolut log2FC≥1.

One of the advantages offered by the nCounter® panels is their sensitivity, as they can be used without prior mRNA extraction. This is intended for cells and, to our knowledge, it has not been employed for EV analyses yet. Considering that EV isolation from tissue already is a multistep process (with the risk of losing material and information at every step of the protocol, we thought it could be of great advantage to bypass the mRNA isolation step, as the latter may result in mRNA loss. Hence, as a proof of concept, we ran a second nCounter® Mouse Neuropathology panel for which the mRNA was not isolated before (from now onwards termed “NI (non-isolated) samples” versus “I (isolated) samples”). The heat map and volcano plot in Fig. 2C and 2D show that from the “NI”-samples, we obtained several significantly upregulated mRNAs for the tMCAO BDEVs compared to shams (88 mRNAs upregulated with a log2FC ≥1 and 11 mRNAs downregulated with log2FC ≤-1; Fig. 2C, Table 1 and Suppl. Table 2). With some differences in the fold-change, “I” and “NI” samples shared most of the highly upregulated mRNAs. Thus, out of the significantly upregulated mRNAs (log2FC≥2) in tMCAO (31 in the “I”-samples panel, and 29 in the NI-samples panel), 23 were common (Table 1 and Venn diagram in Fig. 2E). Eight mRNAs (Tgfb1, Tspo, Cxcl16, Grn, Ccr2, Hpgds, Itga5, and Spi1) present in the isolated samples panel with a log2FC≥2 showed a lower fold-change in the “NI” samples panel, although still showing upregulation (log2FC≥1). Vice-versa, five mRNAs (Lrrc25, Tnfrsf1b, Bcas1, Tnfrsf1a, and Irf8) showed a log2FC≥2 in the “NI” samples, which was lower (although still upregulated with a log2FC≥ 1) in the “I” samples (Table 1). “I” and “NI” samples also shared most of the differentially expressed mRNAs (76 mRNAs), meaning that 81% of the absolute log2FC≥1 mRNA significantly differentially expressed in the “I”-samples are likewise significantly differentially expressed in the “NI”-samples. Because differences between the panels could be attributed to panel batches or sample variation in the tMCAO model, we conclude that overall the nCounter® panels are suitable for the study of BDEVs mRNA cargo without the need for prior mRNA extraction.

**Table 1.**
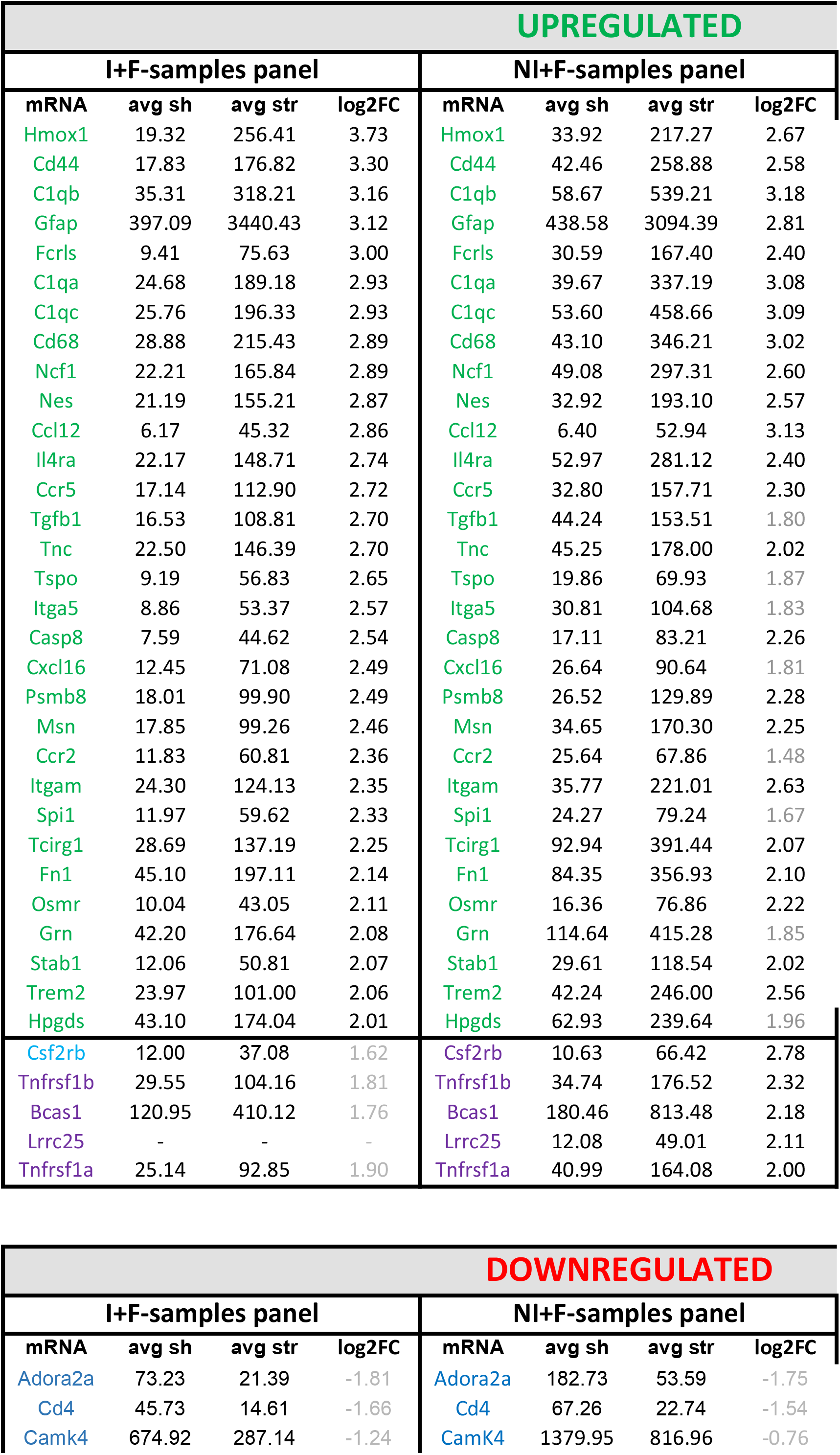
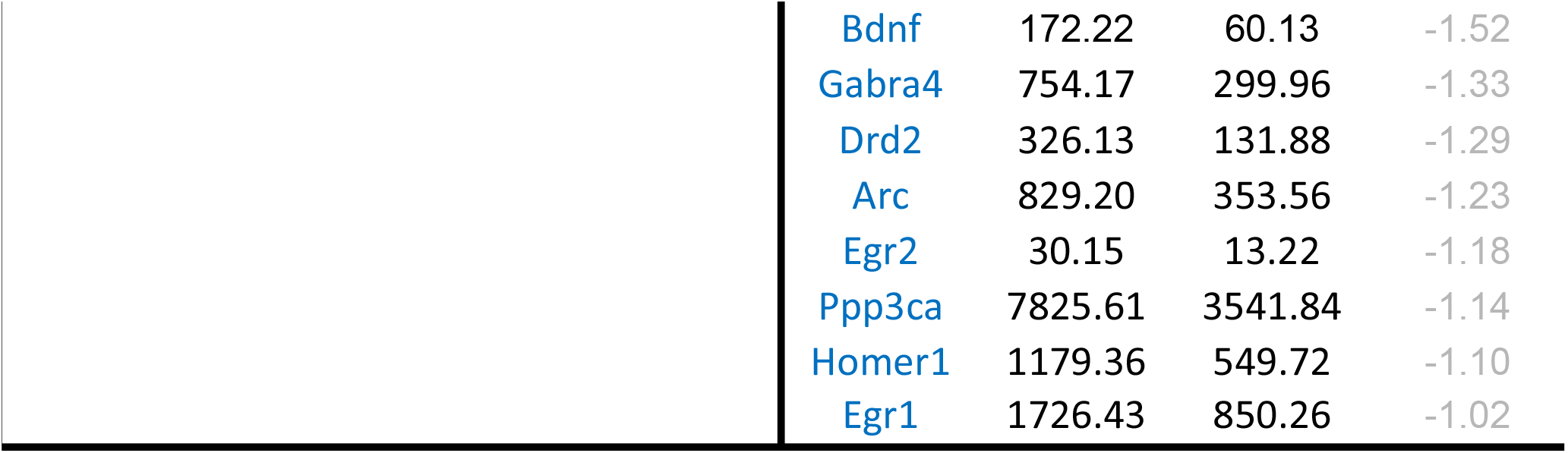
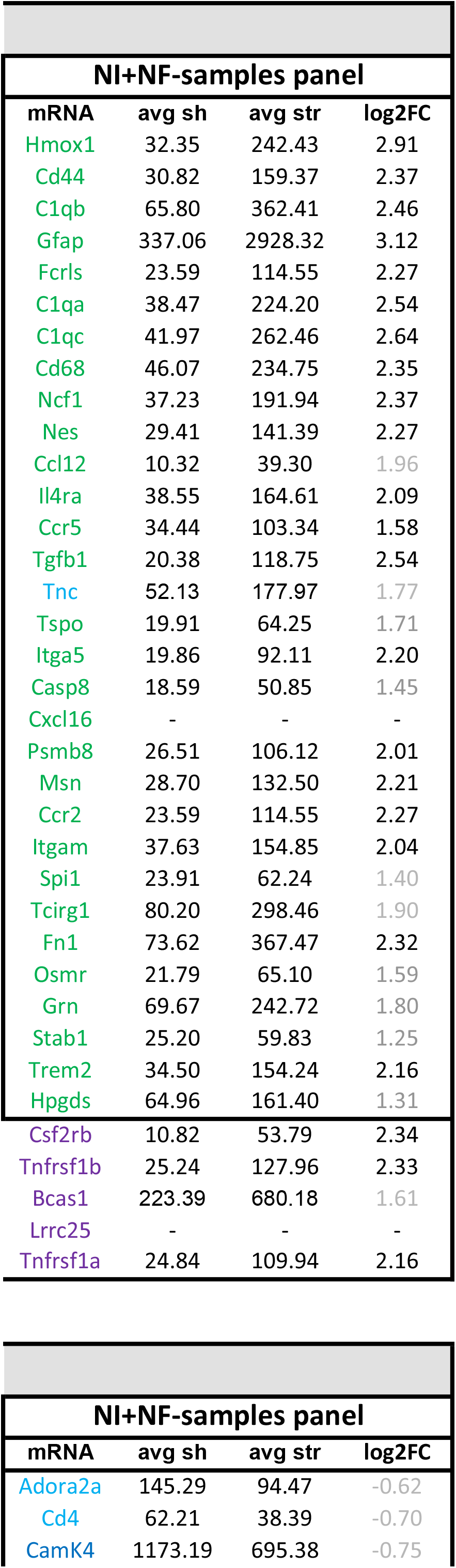
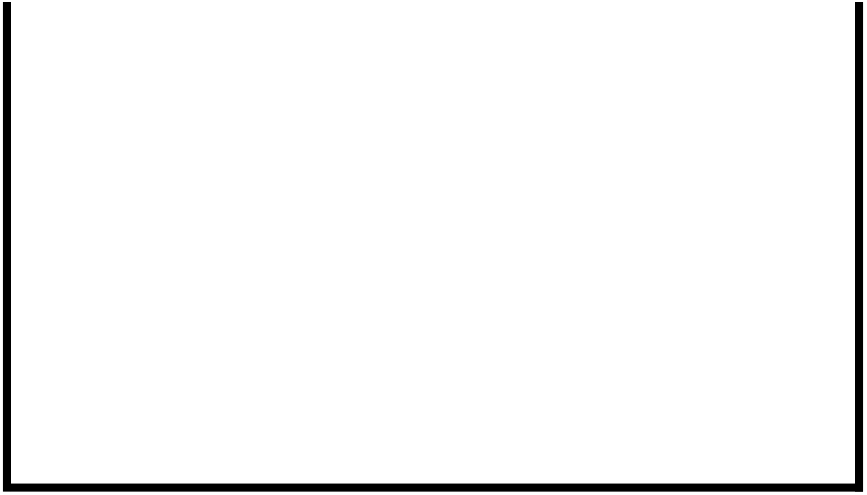
Comparison between upregulated (log2FC≥2) and downregulated mRNAs in isolated vs non-isolated and non-filtered mRNAs from BDEVs in tMCAO compared to shams. Mean counts, and log2FC for each group are provided. Samples with isolated mRNAs and a filtration step (column 1, I+F samples) are taken as the paradigm. The mRNAs of the other samples are compared to column 1 (and therefore the FC is not in decreasing order). In light grey is the fold change that is less than 2 for the mRNAs than in the I+F samples have a log2FC≥2. In blue, mRNAs that did not satisfy the criteria of FDR≤0.1 but are included in the list for comparison. In violet, mRNAs that were found increased with a log2FC≥2 in the NI+F samples which are then compared with the other two columns. No specific mRNA was found in the NI+NF samples. Downregulated mRNAs are shown in dark blue. None of the downregulated mRNAs were observed with a log2F ≤-2, and therefore, the FC is in grey. CxCl16 and Lrcc25 are not present in all columns as, in these instances, they did not satisfy the premises stablished for the background.

In a previous study, we have shown that introducing a filtration step with a 0.2µm membrane leads to the differentiation between two EV populations harboring different protein content, with the population with a diameter ≤ 200 µm (i.e., small BDEVs, sBDEVs) being relatively enriched in ribosomal proteins^32^. The results presented above were generated with samples undergoing this filtration step. However, we were also interested in knowing whether the relative mRNAs in BDEV populations would change by skipping this filtration step and including larger EVs as well. Therefore, in the same panel (all “NI”-samples), we also added BDEVs samples that were lacking the 0.2 µm filtration step (“NF” samples) and compared them to the filtrated (“F”) samples described above. As shown in the heat map and the volcano plot (Fig. 3A and 3B) and Table 1, the most upregulated mRNAs (log2FC≥2) present in the “NF” samples were also among the most upregulated present in the filtrated samples, although the relative fold-change slightly varied. In the “NI”+”NF” samples, we found 22 mRNAs upregulated with a log2FC≥2, and 60 mRNAs upregulated when the cut-off was set to log2FC≥1. In these samples, no significantly downregulated mRNA species with a log2FC≤-1 were found (Table 1 and Suppl. Table 3). As shown in the Venn diagrams in Fig. 3C and 3D, no mRNA was exclusively detected when the samples were not filtered. These findings could indicate that the majority of mRNAs are contained in small EVs (≤ 200 µm). Alternatively, our observations may suggest that because bigger EVs have a relatively higher protein content, the background of the panels is increased thereby relatively lowering mRNA detection, which could explain the generally low fold-change detection in 2 out of the 3 samples shown in the heat map (Fig. 3A).

**Figure 3.**
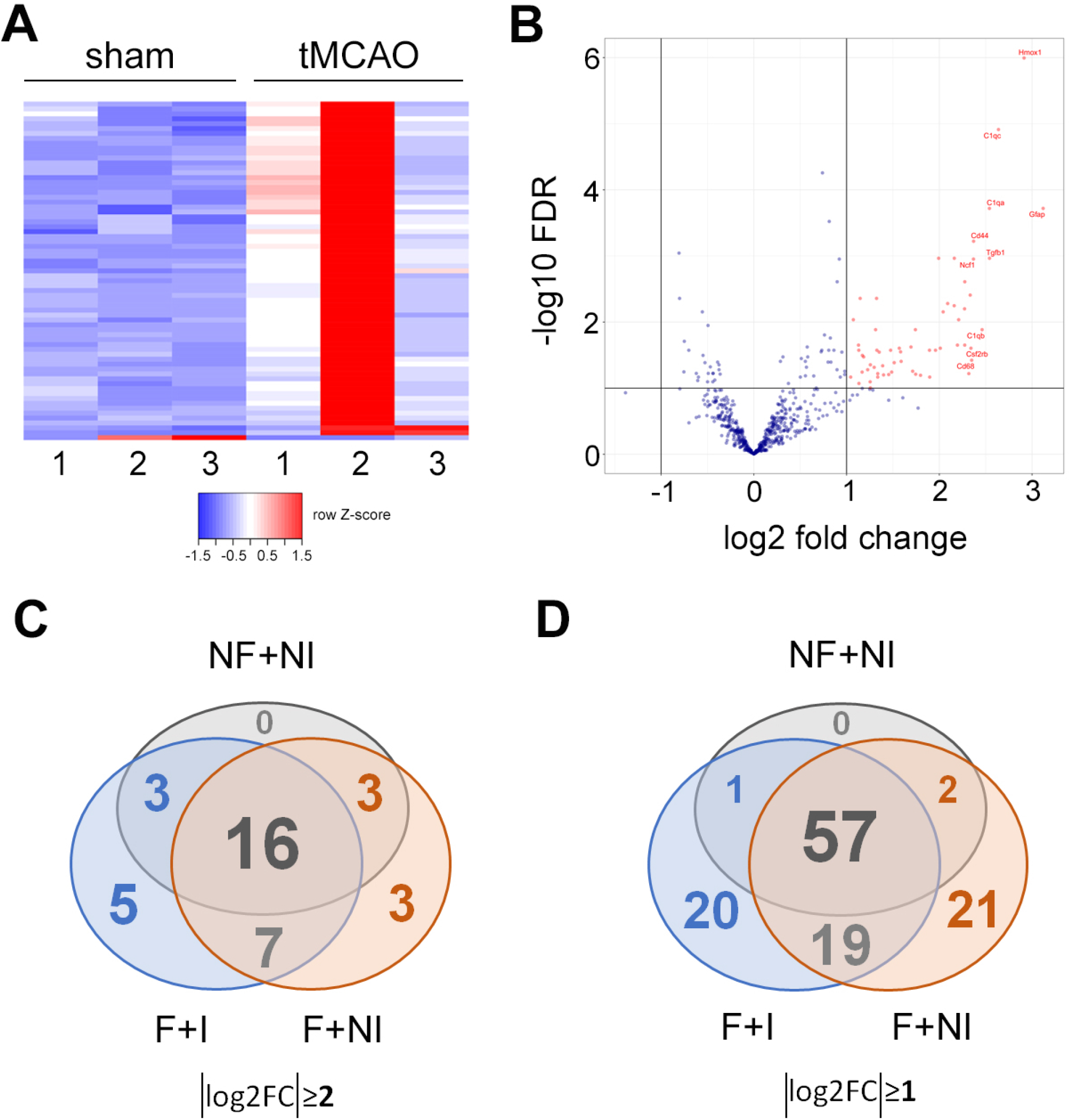
Non-filtered BDEVs show no major differences compared to the filtered ones in mRNA content after tMCAO. (A) Heat map of the up- and downregulated mRNAs (absolut log2FC≥1) in BDEVs of tMCAO compared to shams (n=3 mice per group). (B) Volcano plot for the same mRNAs as in (A) displaying the names of the ten significantly differentially expressed mRNAs in BDEVs from tMCAO mice compared to shams. (C) Overview of the most up and downregulated mRNAs (absolut log2FC≥2) in each panel displayed in a Venn diagram showing the number of the commonly shared and the unique mRNAs for each studied condition: F= BDEV samples were filtrated during preparation (sBDEVs); NF= BDEVs samples were non-filtrated during preparation; I= the mRNA was isolated from BDEVs; NI= the mRNA was not isolated before running the nCounter® panel. (D) Venn diagram for the different sample conditions as in (C) but with a cut-off of absolut log2FC ≥1.

### Upregulated mRNAs in BDEVs after tMCAO are (i) related to inflammatory responses, stress defense, and recovery processes, (ii) are mainly released by microglia, and (iii) top hits present in full-length

To validate the data obtained with the nCounter® panels, we next performed RT-qPCR for 5 of the top mRNAs found upregulated in the tMCAO samples. As shown in Fig.4A, the amount of mRNA extracted from BDEVs was sufficient to perform reverse transcription to cDNA and RT-qPCR analyses (n=7 for each group). All the examined mRNAs (Hmox1, Cd44, C1qa, Gfap, and Fcrls) showed a significant increase in BDEVs from tMCAO mice compared to shams with a similar fold-increase as found in the nCounter® panels (Table 1).

**Figure 4.**
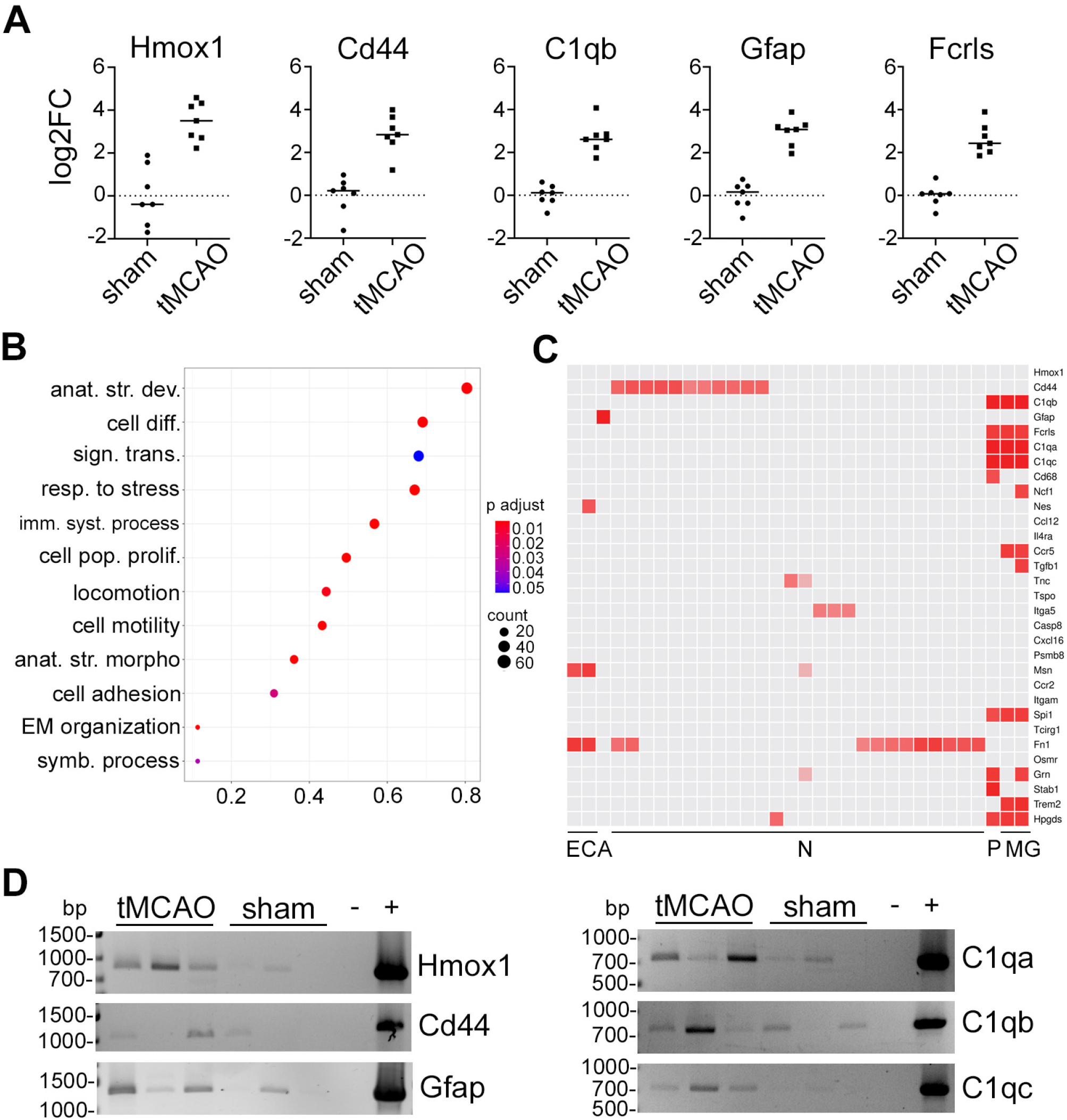
Small BDEVs after tMCAO are enriched in mRNAs related to inflammation, defense response, and recovery processes, mainly derived from microglia, and PCR confirms that sBDEVs contain full-length mRNAs. (A) Dot plots of the RT-qPCR results of 5 of the top upregulated mRNAs in sBDEVs after tMCAO compared to shams. Every dot represents the log2FC calculated from the ΔΔCT value obtained for each mice (n=7 per group). (B) Enriched GO terms dot plot showing the processes related to the most upregulated SBDEV-derived mRNAs (log2FC≥2) in tMCAO compared to shams. The size of the dots is proportional to the number of mRNAs included in each process whereas the color indicates the *p adjusted* value. The list and the full names of the GO terms can be found in Table 2. (C) Heat map adapted from the transcriptomics data of the Allen Brain Atlas. EC: endothelial cells; A: astrocytes; N: Neurons; P: perivascular macrophages; MG: microglia. (D) Agarose gel images of the PCR products for the six most upregulated mRNAs in sBDEVs from tMCAO brains compared to shams. The results are only qualitative as the cDNA used for the PCR was not measured before the PCR (i.e., the cDNA was added to the reaction by volume). Note that all mRNAs tested present a full-length ORF when compared to the cDNA of wild-type mouse brains used as positive controls (+). (-) is the negative control, where all the procedure was performed without cDNA.

Overrepresentation analysis of GO terms by using the 770 genes present in the nCounter® panel as background mRNA, revealed two main types of responses: on the one hand, these were “immune system process”, and “response to stress”, indicating that BDEVs carry mRNAs related to inflammatory processes. On the other hand, terms such as “anatomical structure development”, “cell differentiation”, “cell population differentiation”, “anatomical structure formation involved in morphogenesis”, or “extracellular matrix reorganization”, were also overrepresented, suggesting that certain recovery processes are already taking place at 72 h after tMCAO with BDEVs participating in these events (Fig. 4B and Table 2). To assess the origin of the highest upregulated mRNAs carried in BDEVs after tMCAO (log2FC≥2) we generated a heat map based on data from the Cell Types Database: RNAseq Data from the Allen Brain Map. This revealed that several mRNAs (13 out of 21 with an assigned cell type) were originating from either microglia or perivascular macrophages (Fig. 4C).

**Table 2.** List of GO terms.

| GO term |  |
| --- | --- |
| GO:0048856 | Anatomical structure development |
| GO:0006950 | Response to stress |
| GO:0030154 | Cell differentiation |
| GO:0007165 | Signal transduction |
| GO:0002376 | Immune system process |
| GO:0008283 | Cell population proliferation |
| GO:0040011 | Locomotion |
| GO:0048870 | Cell motility |
| GO:0048646 | Anatomical structure formation involved in morphogenesis |
| GO:0007155 | Cell adhesion |
| GO:0030198 | Extracellular matrix organization |
| GO:0044403 | Symbiotic process |

Finally, we designed PCR primers to qualitatively assess whether the mRNA of six of the top ten candidates present in BDEVs was full length. We used primers allowing for the assessment of the open reading frame (ORF) as an indicator of intact mRNAs. Although the amounts were very variable between the samples, mRNAs for Hmox1, Gfap, C1qa, C1qb, and C1qc, were all present in BDEVs as full-length based on their expected size (Fig. 4D).

### Oligodendrocytic-EVs are upregulated in the total BDEVs pool at 72 h after tMCAO

We have demonstrated earlier that the main EVs population contributing to the total BDEVs pool under physiological conditions originated from microglia. This situation changed 24 h after tMCAO when astrocytic EVs were found significantly upregulated^32^. Considering our aforementioned finding of a predominantly microglial origin of mRNAs in BDEVs obtained at 72 h after stroke (Fig. 4C), we wondered whether this would also reflect an increased contribution of this cell type to the total EV pool in stroked brains at this time-point. Therefore, by applying the same methodology as in our previous study (i.e., biochemical assessment of two typical markers per brain cell type), we compared BDEVs from tMCAO (n=5) and sham (n=5) mice at 72 h after stroke. For neurons, we detected the Neural Cell Adhesion Molecule 1 (NCAM1) as it was present in the mass spectroscopy analysis from BDEVs in our previous study, and Synapsin 1 as used before^32^. As astrocyte-specific markers we assessed the Excitatory Amino Acids Transporter 1 and 2 (EAAT1, EAAT2)^39^; and for microglia, we used the G-protein coupled P2Y receptor 12 (P2Y12) as well as CD40, present in activated microglia and macrophages^40^. Lastly, for oligodendrocytes, we detected the proteolipid protein (PLP, a major component of the myelin sheath) and 2′-3′-Cyclicnucleotide 3′-phosphodiesterase (CNP) ^41^. The quantification was done by first referring the band intensity of the marker protein to the total protein staining serving a loading control, and the mean values between sham and tMCAOs were then compared. As shown in the western blots of Fig. 5A, all marker proteins were present in BDEVs. An obvious increase in both markers for oligodendrocytes (i.e., PLP and CNP1, significantly upregulated in tMCAO \*\**p*=0.0079 and \**p*=0.0317, respectively) was observed. This increase was specific for BDEVs as brain homogenates showed no differences between shams and strokes (Suppl. Fig. 2). Surprisingly, although most of the mRNA present in BDEVs in our study is characteristic of microglial origin, both microglial markers used for immunoblotting did not show any increase in the tMCAO samples compared to shams. The neuronal markers NCAM1 and Syn1 were upregulated but only the increase in NCAM1 reached significance (\*\**p*=0.0079), indicating that neurons could significantly contribute to the BDEVs pool 72 h after stroke. Again, as for PLP and CNP1, the increase in NCAM was specific for BDEVs and not paralleled in brain homogenates (Suppl. Fig. 2). Finally, although Gfap was one of the mRNAs with highest fold-change in our analysis, we did not observe a significant increase in BDEVs from astrocytes. This is in contrast to what we have previously found at 24 h after tMCAO using the same protein markers^32^ and may highlight the transient nature of successive inflammatory and repair processes after stroke with the relative contributions from different cell types changing over time.

**Figure 5.**
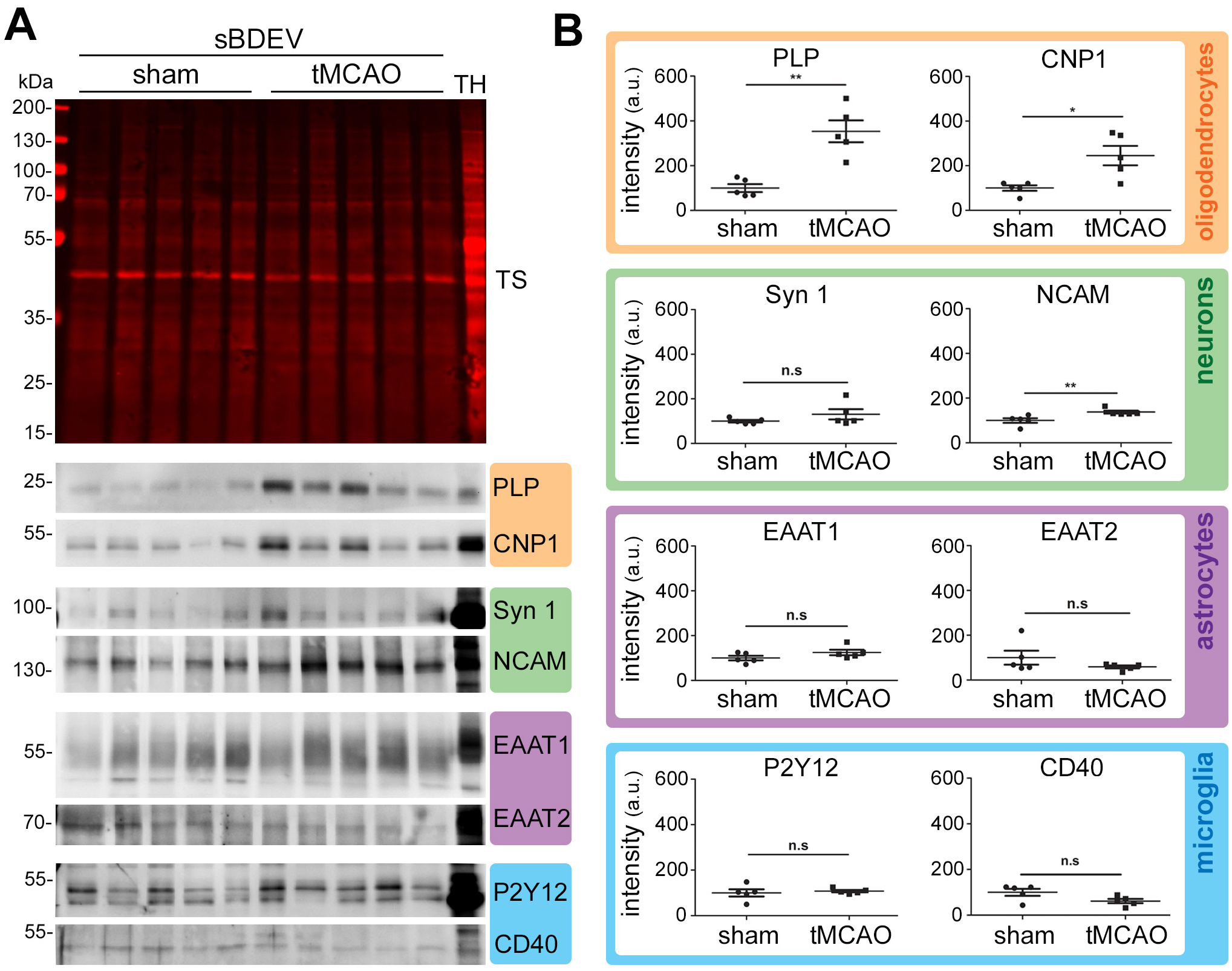
The contribution of oligodendrocytes to the BDEVs pool is significantly upregulated at 72 h after tMCAO. (A) Western blots of sBDEVs samples from tMCAO and shams (n=5 per group) blotted for cell-type-specific markers: PLP and CNP1 are used as protein markers for oligodendrocytes (orange frame); synapsin1 (Syn1) and NCAM as markers for neurons (green); EAAT1 and EAAT2 as protein markers for astrocytes (pink) and P2Y12 and CD40 as markers for microglia/macrophages (blue). TH is a total mouse brain homogenate loaded in parallel for comparison purposes. TS is a representative total protein staining of the nitrocellulose membranes (TSs of all blots used for these analyses are provided in Suppl. Fig. 3). (B) Quantifications of the western blot intensities. For the quantification, each band intensity was first referred to the corresponding lanes of the total protein staining. Both markers for oligodendrocytes were found significantly increased upon stroke. Regarding neuronal markers, NCAM was significantly increased while Syn1 only showed a tendency to be elevated. Exact *p-* values are given in the main text.

**Figure 6.**
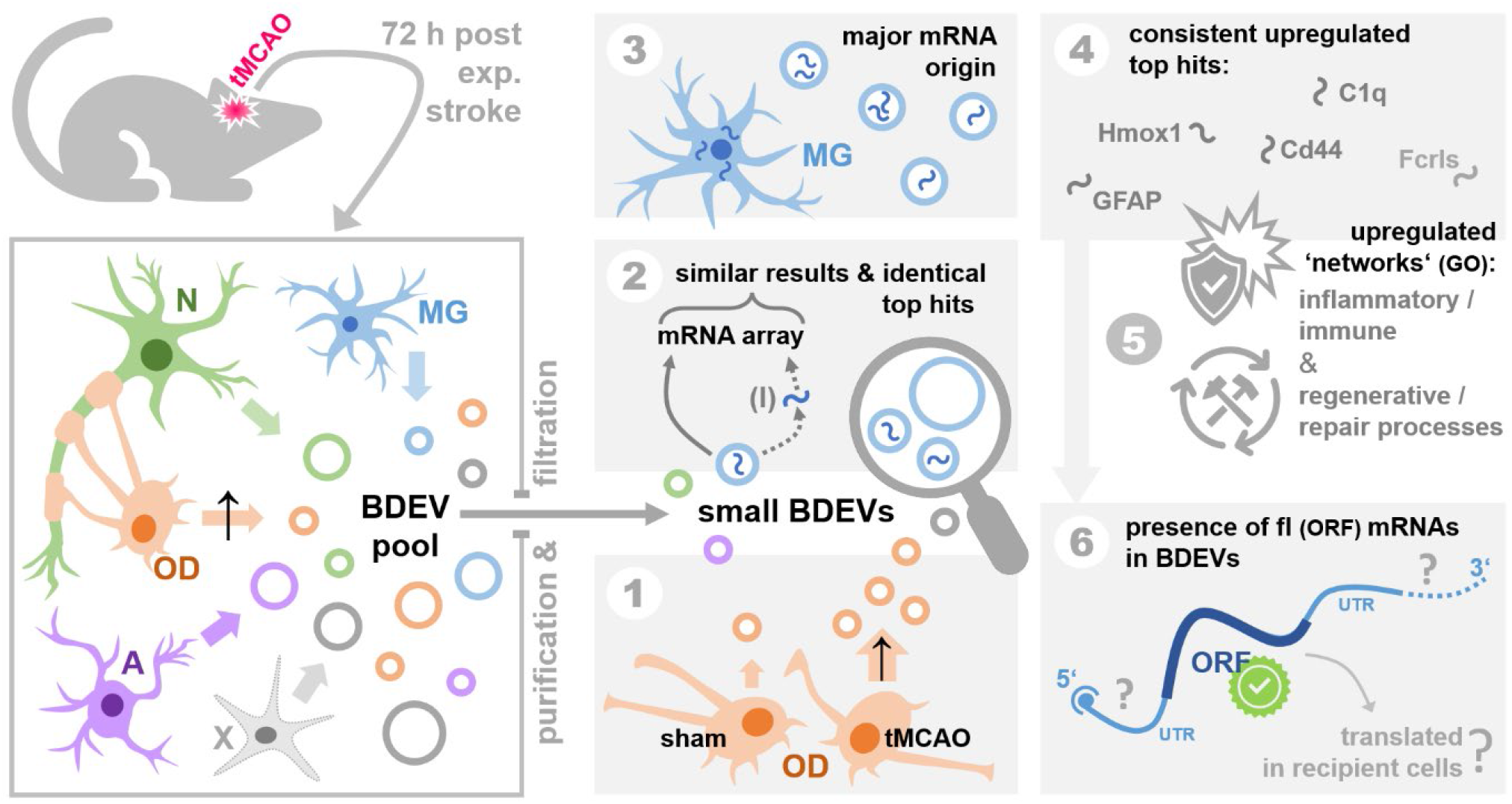
Summarizing scheme of the main findings and open questions. This study assessed alterations in the brain EV pool and the mRNA composition of BDEVs 72 hours after experimental stroke (tMCAO) and reperfusion in mice. Neurons (N), microglia (MG), oligodendrocytes (OD), astrocytes (A), and several other cell types (X) contribute to the EV pool in brain (box on the left). Focussing on small BDEVs, we found an upregulated (↑) contribution by ODs to the latter at this time point after stroke (①). Comparing samples with (I) and without RNA isolation, our multiplexed mRNA arrays revealed similar results regarding mRNA upregulated candidates, and our comparison of filtered and non-filtered BDEVs indicated that mRNAs may be predominantly contained in small BDEVs (②). Though OD showed an increased contribution to the overall EV pool, mRNAs assessed in BDEVs were mainly of MG origin (③). Consistent top hits showing an increased abundance in BDEVs after stroke included mRNAs for Hmox1, Gfap, Cd44, C1q proteins, and Fcrls (④), and predominant GO terms derived from our mRNA findings could largely be linked with the aspects of inflammatory/immune and regenerative/repair regulation (⑤), thus fitting to processes expected to be initiated and to take place at this time point after stroke. Notably, we confirmed the presence of ‘intact’ mRNAs (as judged by a full-length (fl) ORF) for 6 top candidates upregulated in BDEVs upon stroke (⑥). Whether these mRNAs also contain important elements such as the 5’ cap structure or the 3’ poly-A tail and, importantly, if these mRNAs are successfully taken up and possibly even translated in recipient cells to elicit (fast) biological responses (e.g., in the context of post-stroke regeneration) remains to be studied further.

## DISCUSSION

In the present study, we have used the Nanostring nCounter® panels to demonstrate that, after inducing stroke in mice followed by 72 h reperfusion, BDEVs increased their content in mRNAs related to immune, inflammatory, and defense responses as well as recovery processes. The mRNAs present a full-length ORF in BDEVs, at least for the most upregulated genes in tMCAO compared to shams. The top five most upregulated genes shared by all panels detected here are Hmox1, Cd44, C1q (composed by C1qa, C1qb, and C1qc), Gfap, and Fcrls. Hmox1 encodes for the inducible Heme Oxygenase (HO1), which has been implicated in neuroprotection after stroke and is considered a possible therapeutic target^42^. CD44 protein is important for synaptic plasticity and axon guidance, among other functions, in the central nervous system (CNS). Under pathological conditions such as multiple sclerosis (MS) and its respective experimental model in rodents (Experimental Autoimmune Encephalomyelitis, EAE), CD44 has immunomodulatory properties and protects from the breakage of the BBB^43^. After stroke, CD44 seems to be upregulated in neural stem/progenitor cells (NSPCs) and microglia/macrophages at the penumbra area^44^. C1q, the recognition molecule complex that initiates the classical pathway of the complement cascade, is secreted by macrophages and exerts important functions in the brain such as tagging unwanted synapses for elimination^45,46^. In stroke, early activation of the complement system leads to inflammation, but at later time-points, C1q might also play a role in regenerative processes^47^. Gfap encodes for the Glial Fibrillary Acidic Protein (GFAP) which is mainly expressed by astrocytes. Its expression increases when astrocytes become reactive and when they form the glial scar which limits inflammation and promotes repair^48-50^. Fcrls encodes for the Fc Receptor-like S, a scavenger receptor expressed specifically by microglia^51^ but only in mice, rats, and dogs^52^. Scavenger receptors bind ligands that are non-self or self-altered molecules and remove them by phagocytosis (among other mechanisms) leading to the elimination of degraded or harmful substances^53^. Thus, at 72 h after tMCAO, BDEVs contain mRNAs encoding for proteins involved in inflammatory and defense responses but which may also participate in the reconstruction and repair of the affected area.

Because microglia and astrocytes are reactive and proliferate upon stroke, one question is whether an increase of mRNAs such as Gfap or C1qs in BDEVs is just a consequence of the increased amount of these mRNAs in the parental cells. While with the present study we cannot rule out this possibility (as one would have to assess the amount of mRNA in the cells of origin), the fact that Hmox1, C1q, Cd44, and Fcrls showed in general rather low counts in BDEVs from tMCAO mice (although with a significant fold increase compared to sham), our findings would speak in favor of a specific loading of these mRNAs into EVs as a reaction to hypoxic ischemia.

It has been shown that different types of EVs contain different RNA profiles, for instance, with apoptotic bodies being richer in ribosomal RNA (rRNA) when compared to microvesicles and exosomes^54^. Interestingly, we did not detect major differences in relative amounts of mRNAs between filtered and non-filtered samples (the latter presenting a larger average diameter^32^). Although we cannot rule out that an increased amount of protein (since the RNA was not isolated, and bigger EVs probably contain relatively more amount of proteins) could increase the background (overall fewer mRNAs were detected compared to the “NI+F” panel), it could also imply that sEVs (≤ 200nm) carry the majority of the EV-associated mRNAs. Crescitelli *et al*. showed that in two out of three investigated cell lines, most of the RNA was present in exosomes (which fall into the category of sEVs)^54^. Although the type of sample and the isolation methods impact the RNA yield and size distribution^55^, and though the protocols differ between that and our study, the finding could indicate a general lack (or at least relatively reduced presence) of mRNA in larger EVs. Notably, in our previous study^32^ we showed that sBDEVs were enriched in ribosomal proteins when compared to BDEVs that were not filtered and, in fact, ribosomal proteins are commonly found in BDEVs even when using different isolation protocols^56^. Even though it has been shown that EVs do not have the appropriate machinery to translate mRNA directly within the vesicles (at least in cultured cell lines^10^), it could be hypothesized that they could deliver most of the translation machinery plus the mRNA onto recipient cells, particularly as tRNA have also been found in BDEVs^57^.

Previously, we observed a significant increase in astrocytic BDEVs in the total brain pool of tMCAO mice compared to shams at 24 h after stroke^32^. In the present study investigating the time-point of 72 h after tMCAO, EVs from oligodendrocytes are the most upregulated species as judged by two protein markers in western blot analyses. This came as a surprise considering that much of the BDEVs mRNA was found to be significantly upregulated at this time-point after tMCAO could be ascribed to microglia. However, given the fact that our analyses only investigated relative changes upon stroke and not absolute contributions by cell types, both findings are not necessarily conflictive as microglia derived-EVs could still make up a large part of the total pool. Of note, oligodendrocytes, the myelinating cells of the CNS, play a currently understudied role in stroke. They are highly susceptible to ischemic conditions, and the remyelination process starts through the proliferation of oligodendrocyte progenitor cells (OPCs) within the penumbra area^58^. Apart from axonal myelination oligodendrocytes can have an influence on neuronal survival after stroke. For instance, *in vitro* studies show that upon certain neuronal signals, oligodendrocytes release EVs that rescue primary neurons subjected to oxygen-glucose deprivation (OGD, an *in vitro* model of stroke)^29,30^. Oligodendrocyte-derived EVs also influence axonal transport in physiological conditions, an effect that is enhanced when neurons are nutrient-deprived, helping neurons to survive through the delivery of stress-protective macromolecules such as heat shock proteins^28^. Microglia-to-oligodendrocyte communication is another axis that should be considered. Oligodendrocytes can release chemokines and other factors (presumably through EVs) that, under stress conditions, promote microglial phagocytosis^59^. In a model of demyelination in mice, microglia and macrophages turn to anti-inflammatory/immunoregulatory (M2) states and promote the differentiation of OPCs into oligodendrocytes^60^. Thus, by the fact that we also observe an increase in oligodendrocytic-derived EVs at 72 h after tMCAO (a time-point when recovery processes start to take place), it seems plausible that EVs of this particular origin not only play a role in the neuronal remyelination after injury but also in supporting neuronal survival and debris clearance, possibly through communication with microglia. From a technical point of view, we here present that the Nanostring nCounter® panels are useful tools to study the mRNA composition in BDEVs. We chose this technology because while RNA-seq is a very sensitive method and can detect a wide range of RNAs in a given sample, the nCounter® technology offers the advantage that the starting material can be of low amounts and that no further steps (such as reverse transcription, which can introduce certain biases) are necessary to analyze the mRNA content^61^. We show that even when the RNA was not extracted from BDEVs, similar results (although differing in absolute amounts) were obtained compared to samples that underwent the additional RNA isolation step. This is a clear advantage, as one of the problems when assessing EV-derived mRNA, and a considerable challenge for this field is the use of different methodologies to isolate EVs’
s RNA^62^. Moreover, the approach presented here reduces steps in sample preparation and probably increases the chances to detect some species that otherwise would be lost during mRNA isolation. In our case, we were able to detect more significantly downregulated RNA species. It has recently been described that nCounter® panels are also useful and sensitive to analyze EVs isolated from human plasma samples and supernatants of cell cultures. However, in these studies, the amount of EVs obtained was too low to run the samples directly, and intermediate steps such as RNA retro-transcription and cDNA amplification had to be introduced^63,64^. These studies confirmed that even when RNA extracted from EVs is present in low amounts, it can still be used for analysis with the nCounter® panel. Our present report on BDEVs, to the best of our knowledge, is the first to show the suitability of these arrays for analysing the mRNA content of EVs obtained from complex tissues such as brain.

Some papers described the presence of mRNA in exosomes that could be translated in recipient cells^10,14,65^, while in other instances mRNA in EVs was found mainly to be fragmented, as in the case of EVs from human glioblastoma stem cells^3,66^. A recent study using EVs isolated from human blood and performing RNAseq, found that a substantial amount of the total RNA detected in EVs was full-length with a mean size of around 2.800 nucleotides, but also containing very large mRNAs (e.g., KMT2D with more than 19.000 bp)^67^. In our study, we have explicitly checked by PCR if we could amplify the full open reading frame (ORF) of six of the top upregulated candidates. We confirmed that they were all present as full-length in BDEVs, and therefore have at least the potential to be translated in the recipient cells. However, studying the biological relevance of the mRNA content (and other RNA cargoes) in EVs and proving their functionality in the recipient cells is very challenging^68^. It has been shown in EVs of human glioblastoma stem cells that the number of mRNAs is substantially low, as it was estimated to be 1 copy in 1,000 EVs for the most abundant mRNA, and for others less than 1 copy every 10,000 to 100,000, or even 1 copy in a million EVs for some RNA species. However, related to miRNA and despite the low numbers, the authors of this study observed an effect of miR21 in recipient cells^3^. We have also made some rough calculations on the possible number of EVs containing mRNA detected in our system: for the NI-samples (where, in general, the amounts of mRNA counts were higher), we loaded 2.8 µg of sample which approximately corresponds to 6.6×10^10^ EVs (calculation based on the mean of 6 NTA measurements and previous micro-BCA assays). If we take Gfap (for which we confirmed presence in full length, and which showed the highest counts among the significantly upregulated BDEV mRNAs in stroke), the ratio would be approx. 1 mRNA copy per 19,000,000 EVs. Thus, the biological significance of mRNAs present in BDEVs in these relatively low amounts is difficult to judge with the current knowledge and tools. We have attempted to incubate BDEVs with several cell lines to assess any putative transfer and translation of Gfap and Hmox1 in recipient cells by employing immunocytochemistry and confocal microscopy to no avail (data not shown). However, several variables could have played a role in our negative results. In the scenario where EVs were delivered and the mRNA translated, it is plausible to think that either confocal microscopy was not sensitive enough to detect newly translated proteins or the time points we chose to assess protein translation (24 h and 48 h) were not appropriate.

An important question in the EV field is how the cargo is delivered to the cytosol, especially in the case of RNAs that presumably have to reach this compartment in the recipient cell to exert their function/be translated to proteins^68^. Direct fusion of EVs with the target cell’
ss plasma membrane would be the most plausible mechanism to deliver RNA and the EV luminal content to the cytosol of recipient cells. However, in only a few instances fusion of EVs with recipient cells has been demonstrated until now^69-71^, and most of the experiments point to endocytosis as the main take-up mechanism. This implies fusion with late endosomal membranes as a mechanism of endocytic escape for the content of EVs to eventually reach the cytosol^72-74^.

In conclusion, we demonstrate here that nCounter® panels are a useful tool for the targeted study of mRNAs contained in EVs derived from brain tissue. Investigations of the EVs content and alterations therein after cerebral ischemia will surely contribute to increasing our knowledge of this complex pathophysiology and may provide novel therapeutic tools to rescue neurons at the penumbra soon after hypoxic insults.

## ACKNOWLEDGMENTS

The authors would like to thank Prof. Lucie Carrier and Elisabeth Krämer from the Nanostring Core Facility of the UKE for their help and guidance with the nCounter® panels. We also would like to thank Oliver Schnapauff for performing the tMCAO surgery, Ellen Orthey and Marco Lukowiak for helping with the Bioanalyzer and RT-qPCR, and Dr. Christoph König from Nanostring for his valuable help in interpreting the data. A.Bub is a recipient of a scholarship from the “Else Kröner -Promotionskolleg Hamburg – Translationale Entzündungsforschung” (iPRIME). This work was supported by grants from the Werner Otto Stiftung (to Berta Puig) and the Hermann and Lilly Schilling Foundation (to Tim Magnus).

