## Supplemental Figure Legends for "Multiplexed mRNA analysis of brain-derived extracellular vesicles upon experimental stroke in mice reveals increased mRNA content related to inflammation and recovery processes"

**SUPPLEMENTAL TABLES**

**Suppl. Table 1.- List of the mRNAs found differentially upregulated (log2FC≥1) in BDEVs of tMCAO compared to shams when mRNA was isolated.** Downregulated mRNAs are already shown in Table 1.

**Suppl. Table 2.- List of the mRNAs found differentially upregulated (log2FC≥1) in BDEVs of tMCAO compared to shams when mRNA was not isolated.** Downregulated mRNAs are already shown in Table 1.

**Suppl. Table 3.- List of the mRNAs found differentially upregulated (log2FC≥1) in BDEVs of tMCAO compared to shams when mRNA was not isolated and BDEVs were not filtered.** Downregulated mRNAs are already shown in Table 1.

**SUPPLEMENTAL FIGURES**

**Suppl. Fig. 1.- Principal component analysis (PCA) score plots show that BDEVs from shams differentially cluster from BDEVs in tMCAO in all three panels.** The first two principal components are plotted for each sample. Red circles represent the shams whereas blue triangles represent tMCAO samples.

**Suppl. Fig. 2.- No increase in PLP, CNP1, and NCAM in total brain homogenates from tMCAO mice.** (A) Western blots of total brain homogenates from sham and tMCAO samples developed with PLP, CNP1, and NCAM antibodies. TS is total protein staining. (B) Dot plots showing the quantifications for the western blots in (A).

**Suppl. Fig. 3.- Total stainings used for the quantifications shown in Fig.5.** (A) Total staining (TS) for CD40. (B) TS for P2Y12. (C) TS for synapsin, CNP, and PLP. (D) TS for EEAT1 and NCAM. (E) TS for EAAT2. When several antibodies were used for one membrane, the latter was cut according to the molecular weight of the protein of interest, but never stripped and re-incubated again.
