## Supplemental Table 1 for "Multiplexed mRNA analysis of brain-derived extracellular vesicles upon experimental stroke in mice reveals increased mRNA content related to inflammation and recovery processes"

Suppl. Table 1

| mRNA | avg sh | avg str | log2FC | Padj |
|---|---|---|---|---|
| Hmox1 | 19.32 | 256.41 | 3.73 | 8.04E-31 |
| Cd44 | 17.83 | 176.82 | 3.30 | 4.10E-09 |
| C1qb | 35.31 | 318.21 | 3.16 | 1.45E-39 |
| Gfap | 397.09 | 3440.43 | 3.12 | 5.97E-75 |
| Fcrls | 9.41 | 75.63 | 3.00 | 1.15E-11 |
| C1qa | 24.68 | 189.18 | 2.93 | 1.43E-23 |
| C1qc | 25.76 | 196.33 | 2.93 | 2.91E-28 |
| Cd68 | 28.88 | 215.43 | 2.89 | 4.40E-06 |
| Ncf1 | 22.21 | 165.84 | 2.89 | 2.04E-18 |
| Nes | 21.19 | 155.21 | 2.87 | 1.51E-23 |
| Ccl12 | 6.17 | 45.32 | 2.86 | 4.18E-07 |
| Il4ra | 22.17 | 148.71 | 2.74 | 7.72E-18 |
| Ccr5 | 17.14 | 112.90 | 2.72 | 1.64E-15 |
| Tgfb1 | 16.53 | 108.81 | 2.70 | 7.09E-17 |
| Tnc | 22.50 | 146.39 | 2.70 | 3.20E-18 |
| Tspo | 9.19 | 56.83 | 2.65 | 8.63E-09 |
| Itga5 | 8.86 | 53.37 | 2.57 | 3.56E-07 |
| Casp8 | 7.59 | 44.62 | 2.54 | 1.57E-06 |
| Cxcl16 | 12.45 | 71.08 | 2.49 | 1.64E-10 |
| Psmb8 | 18.01 | 99.90 | 2.49 | 1.67E-13 |
| Msn | 17.85 | 99.26 | 2.46 | 2.22E-13 |
| Ccr2 | 11.83 | 60.81 | 2.36 | 5.34E-09 |
| Itgam | 24.30 | 124.13 | 2.35 | 8.70E-17 |
| Spi1 | 11.97 | 59.62 | 2.33 | 6.50E-09 |
| Tcirg1 | 28.69 | 137.19 | 2.25 | 4.87E-11 |
| Fn1 | 45.10 | 197.11 | 2.14 | 9.06E-17 |
| Osmr | 10.04 | 43.05 | 2.11 | 3.50E-05 |
| Grn | 42.20 | 176.64 | 2.08 | 1.02E-13 |
| Stab1 | 12.06 | 50.81 | 2.07 | 5.33E-05 |
| Trem2 | 23.97 | 101.00 | 2.06 | 6.50E-09 |
| Hpgds | 43.10 | 174.04 | 2.01 | 1.72E-14 |
| Trf | 289.47 | 1146.01 | 1.98 | 4.29E-26 |
| Lox | 13.95 | 54.04 | 1.96 | 6.19E-05 |
| Cx3cr1 | 192.01 | 739.88 | 1.94 | 2.37E-24 |
| Cp | 19.72 | 74.71 | 1.92 | 1.08E-06 |
| Eng | 16.49 | 61.07 | 1.90 | 3.78E-07 |
| Tnfrsf1a | 25.14 | 92.85 | 1.90 | 7.14E-10 |
| Gusb | 18.90 | 69.41 | 1.88 | 1.43E-06 |
| Irf8 | 18.80 | 67.70 | 1.84 | 9.74E-07 |
| Stat3 | 65.94 | 233.66 | 1.83 | 4.25E-14 |
| Csf1r | 94.92 | 334.07 | 1.81 | 6.43E-22 |
| Tnfrsf1b | 29.55 | 104.16 | 1.81 | 6.50E-09 |
| Il1r1 | 32.84 | 112.62 | 1.77 | 1.50E-07 |
| Bcas1 | 120.95 | 410.12 | 1.76 | 0.000299 |
| Il6ra | 18.10 | 60.08 | 1.73 | 9.75E-06 |
| Hexb | 138.73 | 462.43 | 1.73 | 1.02E-13 |
| Ccnd1 | 281.07 | 913.97 | 1.70 | 1.42E-16 |
| Tlr2 | 11.79 | 38.04 | 1.69 | 0.000444 |

| | | | | |
|---|---|---|---|---|
| Csf1 | 33.73 | 107.91 | 1.68 | 3.77E-07 |
| Hspb1 | 29.29 | 92.60 | 1.66 | 9.15E-06 |
| Sp100 | 14.09 | 43.33 | 1.60 | 0.001033 |
| Stat1 | 40.98 | 122.48 | 1.59 | 1.04E-06 |
| Plcb3 | 21.17 | 63.39 | 1.58 | 2.22E-05 |
| Nfe2l2 | 77.10 | 227.04 | 1.56 | 5.03E-12 |
| Notch1 | 32.72 | 96.02 | 1.55 | 1.20E-06 |
| Col4a1 | 33.49 | 96.99 | 1.54 | 2.90E-06 |
| Cspg4 | 13.37 | 39.04 | 1.53 | 0.002946 |
| Cxcr4 | 10.95 | 31.38 | 1.53 | 0.009501 |
| Tgfbr2 | 40.57 | 115.81 | 1.51 | 3.34E-08 |
| P2rx7 | 27.01 | 77.53 | 1.51 | 0.041231 |
| Cd9 | 27.78 | 78.57 | 1.49 | 2.67E-05 |
| Phf19 | 10.37 | 28.86 | 1.47 | 0.009891 |
| Gjb1 | 62.17 | 167.85 | 1.43 | 4.98E-09 |
| Ltbr | 27.26 | 72.64 | 1.42 | 1.15E-05 |
| Cdk2 | 15.77 | 41.90 | 1.41 | 0.009501 |
| Myc | 44.15 | 116.57 | 1.40 | 1.57E-06 |
| Mmp14 | 25.99 | 68.87 | 1.40 | 2.98E-05 |
| Tlr4 | 11.40 | 29.77 | 1.39 | 0.014332 |
| Cd33 | 22.77 | 59.69 | 1.38 | 0.000155 |
| Itga7 | 17.82 | 45.70 | 1.37 | 0.00163 |
| Itpr2 | 45.28 | 117.49 | 1.37 | 2.22E-05 |
| Gsn | 99.13 | 253.79 | 1.36 | 3.80E-08 |
| Plxnb3 | 48.63 | 124.19 | 1.35 | 4.36E-05 |
| Pecam1 | 13.69 | 34.42 | 1.31 | 0.012856 |
| Tmem119 | 50.03 | 123.79 | 1.30 | 4.75E-07 |
| Pcna | 200.51 | 492.34 | 1.30 | 3.00E-12 |
| Lif | 11.60 | 28.49 | 1.29 | 0.025807 |
| Il10ra | 31.24 | 75.64 | 1.29 | 6.19E-05 |
| Pfn1 | 688.06 | 1670.34 | 1.28 | 9.87E-16 |
| Il13ra1 | 19.62 | 46.92 | 1.27 | 0.002808 |
| Casp6 | 28.75 | 66.22 | 1.21 | 0.000247 |
| Pla2g4a | 12.96 | 29.94 | 1.20 | 0.022685 |
| Pkn1 | 44.96 | 101.68 | 1.18 | 0.000325 |
| Col4a2 | 35.38 | 80.65 | 1.17 | 0.000344 |
| Casp7 | 19.00 | 41.40 | 1.14 | 0.009778 |
| Sgpl1 | 56.17 | 124.47 | 1.14 | 7.88E-05 |
| Cdkn1a | 76.40 | 168.00 | 1.14 | 1.14E-05 |
| Plekho2 | 83.98 | 181.29 | 1.11 | 2.18E-07 |
| Nlrp3 | 14.45 | 30.21 | 1.06 | 0.050228 |
| Arrb2 | 92.39 | 190.25 | 1.04 | 8.07E-06 |
| Hdac7 | 85.29 | 175.53 | 1.04 | 0.000166 |
| Myrf | 113.40 | 233.46 | 1.04 | 5.00E-05 |
| Fgf2 | 15.60 | 31.67 | 1.02 | 0.066744 |
| Mapkapk2 | 44.99 | 90.50 | 1.00 | 0.000828 |
