## Supplemental Table 2 for "Multiplexed mRNA analysis of brain-derived extracellular vesicles upon experimental stroke in mice reveals increased mRNA content related to inflammation and recovery processes"

**Suppl.Table 2**

| mRNA | avg sh | avg str | log2FC | Padj |
|---|---|---|---|---|
| C1qb | 58.67 | 539.21 | 3.18 | 1.47E-25 |
| Ccl12 | 6.40 | 52.94 | 3.13 | 1.76E-07 |
| C1qc | 53.60 | 458.66 | 3.09 | 1.25E-23 |
| C1qa | 39.67 | 337.19 | 3.08 | 3.39E-23 |
| Cd68 | 43.10 | 346.21 | 3.02 | 4.48E-22 |
| Gfap | 438.58 | 3094.39 | 2.81 | 2.74E-33 |
| Csf2rb | 10.63 | 66.42 | 2.78 | 2.25E-10 |
| Hmox1 | 33.92 | 217.27 | 2.67 | 5.49E-21 |
| Itgam | 35.77 | 221.01 | 2.63 | 7.17E-24 |
| Ncf1 | 49.08 | 297.31 | 2.60 | 1.03E-15 |
| Cd44 | 42.46 | 258.88 | 2.58 | 5.24E-15 |
| Nes | 32.92 | 193.10 | 2.57 | 1.12E-22 |
| Trem2 | 42.24 | 246.00 | 2.56 | 2.89E-22 |
| Fcrls | 30.59 | 167.40 | 2.40 | 1.57E-16 |
| Il4ra | 52.97 | 281.12 | 2.40 | 1.47E-17 |
| Tnfrsf1b | 34.74 | 176.52 | 2.32 | 7.56E-17 |
| Irf8 | 35.74 | 179.69 | 2.30 | 4.55E-12 |
| Ccr5 | 32.80 | 157.71 | 2.30 | 6.50E-16 |
| Psmb8 | 26.52 | 129.89 | 2.28 | 1.90E-14 |
| Casp8 | 17.11 | 83.21 | 2.26 | 6.61E-09 |
| Msn | 34.65 | 170.30 | 2.25 | 2.72E-12 |
| Osmr | 16.36 | 76.86 | 2.22 | 1.54E-09 |
| Bcas1 | 180.46 | 813.48 | 2.18 | 2.52E-12 |
| Lrrc25 | 12.08 | 49.01 | 2.11 | 4.84E-05 |
| Fn1 | 84.35 | 356.93 | 2.10 | 2.82E-14 |
| Tcirg1 | 92.94 | 391.44 | 2.07 | 3.75E-16 |
| Stab1 | 29.61 | 118.54 | 2.02 | 9.20E-09 |
| Tnc | 45.25 | 178.00 | 2.02 | 5.26E-07 |
| Tnfrsf1a | 40.99 | 164.08 | 2.00 | 4.17E-09 |
| Nlrp3 | 12.80 | 49.83 | 1.99 | 1.69E-06 |
| Slc11a1 | 9.04 | 36.79 | 1.99 | 5.25E-05 |
| Hpgds | 62.93 | 239.64 | 1.96 | 5.35E-13 |
| Cx3cr1 | 409.28 | 1547.84 | 1.92 | 7.01E-12 |
| Tspo | 19.86 | 69.93 | 1.87 | 3.58E-07 |
| Grn | 114.64 | 415.28 | 1.85 | 7.29E-08 |
| Cybb | 9.90 | 37.26 | 1.85 | 0.000303 |
| Tlr2 | 11.68 | 41.08 | 1.84 | 7.97E-05 |
| Itga5 | 30.81 | 104.68 | 1.83 | 2.69E-05 |
| Cxcl16 | 26.64 | 90.64 | 1.81 | 2.15E-05 |
| Csf1r | 244.52 | 851.83 | 1.81 | 2.16E-12 |
| Tgfb1 | 44.24 | 153.51 | 1.80 | 1.00E-08 |
| Icam1 | 9.43 | 32.81 | 1.79 | 0.000971 |
| Cdk2 | 22.61 | 78.32 | 1.78 | 1.90E-05 |
| Il10ra | 51.64 | 168.11 | 1.69 | 4.44E-07 |
| Hspb1 | 43.41 | 139.81 | 1.67 | 0.001139 |
| Spi1 | 24.27 | 79.24 | 1.67 | 8.42E-05 |
| Cspg4 | 32.60 | 101.55 | 1.66 | 6.37E-07 |
| Hexb | 290.29 | 892.95 | 1.62 | 5.12E-12 |

| | | | | |
|---|---|---|---|---|
| Csf1 | 73.20 | 221.98 | 1.60 | 2.82E-08 |
| Trf | 641.55 | 1918.99 | 1.58 | 1.62E-09 |
| Sp100 | 24.91 | 72.37 | 1.55 | 0.000173 |
| Myc | 64.75 | 187.60 | 1.54 | 4.00E-09 |
| Cd9 | 53.16 | 152.56 | 1.53 | 2.06E-08 |
| Lif | 14.76 | 41.76 | 1.50 | 0.000949 |
| Ccnd1 | 612.13 | 1712.53 | 1.48 | 5.58E-09 |
| Ccr2 | 25.64 | 67.86 | 1.48 | 0.000502 |
| Tgfbr2 | 56.67 | 155.89 | 1.45 | 2.67E-05 |
| Tnf | 10.49 | 29.53 | 1.45 | 0.010063 |
| Ltbr | 34.57 | 97.37 | 1.44 | 6.77E-05 |
| Nfe2l2 | 149.76 | 406.26 | 1.44 | 6.81E-08 |
| Tnfrsf12a | 24.87 | 68.94 | 1.44 | 0.00022 |
| Cd33 | 36.76 | 97.68 | 1.43 | 7.46E-05 |
| Cp | 36.54 | 94.97 | 1.39 | 1.08E-05 |
| Gjb1 | 107.80 | 275.98 | 1.37 | 3.01E-08 |
| P2rx7 | 66.87 | 170.64 | 1.35 | 2.52E-05 |
| Gsn | 132.49 | 336.21 | 1.34 | 1.53E-07 |
| Tmem119 | 94.77 | 237.29 | 1.33 | 7.58E-06 |
| Epha2 | 11.94 | 29.14 | 1.33 | 0.015798 |
| Plcb2 | 29.12 | 73.57 | 1.30 | 0.000737 |
| Gpr84 | 9.84 | 23.14 | 1.29 | 0.043378 |
| Tlr4 | 20.42 | 50.44 | 1.28 | 0.013009 |
| Stat3 | 158.14 | 380.69 | 1.27 | 7.60E-06 |
| Casp7 | 26.94 | 64.04 | 1.23 | 0.0496 |
| Gusb | 57.42 | 133.49 | 1.21 | 7.97E-05 |
| Tnfrsf10b | 10.63 | 22.55 | 1.19 | 0.086192 |
| Psmb9 | 23.98 | 54.63 | 1.18 | 0.005509 |
| Plcb3 | 51.73 | 118.85 | 1.17 | 6.39E-05 |
| Eng | 53.29 | 116.38 | 1.17 | 0.002662 |
| Il1r1 | 76.98 | 171.10 | 1.17 | 6.88E-07 |
| Stat1 | 86.53 | 191.90 | 1.16 | 3.38E-05 |
| Itpr2 | 132.97 | 288.19 | 1.13 | 5.51E-06 |
| Phf19 | 16.22 | 34.71 | 1.09 | 0.047613 |
| Casp1 | 12.67 | 26.97 | 1.09 | 0.053016 |
| Notch1 | 125.95 | 266.27 | 1.09 | 4.81E-05 |
| Il6ra | 45.97 | 96.91 | 1.06 | 0.000442 |
| Pfn1 | 947.83 | 1957.05 | 1.05 | 0.000122 |
| Cdkn1a | 158.06 | 329.32 | 1.05 | 0.000377 |
| Gdnf | 12.95 | 26.17 | 1.04 | 0.091634 |
