## Supplemental Table 3 for "Multiplexed mRNA analysis of brain-derived extracellular vesicles upon experimental stroke in mice reveals increased mRNA content related to inflammation and recovery processes"

**Suppl.Table 3**

| mRNA | avg sh | avg str | log2FC | Padj |
|---|---|---|---|---|
| Gfap | 337.06 | 2928.32 | 3.12 | 0.000191 |
| Hmox1 | 32.35 | 242.43 | 2.91 | 1.01E-06 |
| C1qc | 41.97 | 262.46 | 2.64 | 1.22E-05 |
| Tgfb1 | 20.38 | 118.75 | 2.54 | 0.001084 |
| C1qa | 38.47 | 224.20 | 2.54 | 0.000191 |
| C1qb | 65.80 | 362.41 | 2.46 | 0.013036 |
| Cd44 | 30.82 | 159.37 | 2.37 | 0.000599 |
| Ncf1 | 37.23 | 191.94 | 2.37 | 0.001115 |
| Cd68 | 46.07 | 234.75 | 2.35 | 0.037777 |
| Csf2rb | 10.82 | 53.79 | 2.34 | 0.024867 |
| Tnfrsf1b | 25.24 | 127.96 | 2.33 | 0.003912 |
| Fn1 | 73.62 | 367.47 | 2.32 | 0.060245 |
| Fcrls | 23.59 | 114.55 | 2.27 | 0.002458 |
| Ccr2 | 14.73 | 71.13 | 2.27 | 0.022323 |
| Nes | 29.41 | 141.39 | 2.27 | 0.006308 |
| Msn | 28.70 | 132.50 | 2.21 | 0.00922 |
| Itga5 | 19.86 | 92.11 | 2.20 | 0.022323 |
| Trem2 | 34.50 | 154.24 | 2.16 | 0.001084 |
| Tnfrsf1a | 24.84 | 109.94 | 2.16 | 0.005658 |
| Il4ra | 38.55 | 164.61 | 2.09 | 0.005245 |
| Itgam | 37.63 | 154.85 | 2.04 | 0.00701 |
| Psmb8 | 26.51 | 106.12 | 2.01 | 0.024867 |
| Irf8 | 29.37 | 117.03 | 1.99 | 0.001084 |
| Ccl12 | 10.32 | 39.30 | 1.96 | 0.026683 |
| Tcirg1 | 80.20 | 298.46 | 1.90 | 0.067986 |
| Lif | 11.82 | 43.50 | 1.89 | 0.026683 |
| Grn | 69.67 | 242.72 | 1.80 | 0.063853 |
| Sp100 | 20.36 | 67.93 | 1.74 | 0.061507 |
| Il10ra | 37.08 | 124.34 | 1.74 | 0.013036 |
| Tspo | 19.91 | 64.25 | 1.71 | 0.055405 |
| Trf | 575.98 | 1864.24 | 1.69 | 0.023845 |
| Osmr | 21.79 | 65.10 | 1.59 | 0.039337 |
| Ccr5 | 34.44 | 103.34 | 1.58 | 0.028206 |
| Cx3cr1 | 351.67 | 1042.06 | 1.57 | 0.024867 |
| Nlrp3 | 15.75 | 44.19 | 1.51 | 0.041155 |
| Ltbr | 27.97 | 77.55 | 1.49 | 0.05714 |
| Csf1r | 223.47 | 621.06 | 1.47 | 0.026683 |
| Casp8 | 18.59 | 50.85 | 1.45 | 0.062259 |
| Spi1 | 23.91 | 62.24 | 1.40 | 0.063043 |
| Cd9 | 44.38 | 115.45 | 1.39 | 0.045874 |
| Il13ra1 | 21.80 | 54.77 | 1.35 | 0.028478 |
| Tnfrsf12a | 16.65 | 41.78 | 1.34 | 0.060245 |
| Hexb | 269.80 | 680.96 | 1.34 | 0.067986 |
| Hspb1 | 36.56 | 91.40 | 1.33 | 0.026683 |
| Csf1 | 82.64 | 207.29 | 1.32 | 0.004387 |
| Hpgds | 64.96 | 161.40 | 1.31 | 0.013036 |
| Tgfbr2 | 52.52 | 129.24 | 1.30 | 0.048391 |
| Tlr4 | 13.71 | 32.96 | 1.26 | 0.099062 |

| | | | | |
|---|---|---|---|---|
| Cd33 | 30.02 | 72.73 | 1.26 | 0.044327 |
| Stab1 | 25.20 | 59.83 | 1.25 | 0.080157 |
| Gusb | 42.88 | 100.52 | 1.23 | 0.052605 |
| Myc | 72.28 | 165.01 | 1.18 | 0.033679 |
| Stat1 | 80.47 | 181.63 | 1.17 | 0.03192 |
| Cp | 38.55 | 87.33 | 1.17 | 0.05714 |
| Gsn | 134.99 | 298.80 | 1.15 | 0.004387 |
| Il6ra | 40.85 | 89.43 | 1.13 | 0.084853 |
| Il1r1 | 66.94 | 147.33 | 1.13 | 0.026683 |
| Gjb1 | 86.33 | 188.23 | 1.13 | 0.022323 |
| Stat3 | 130.44 | 274.51 | 1.07 | 0.00922 |
| Tmem119 | 70.78 | 145.42 | 1.04 | 0.067986 |
