## Supplementary figures and images for "Multiplexed mRNA analysis of brain-derived extracellular vesicles upon experimental stroke in mice reveals increased mRNA content related to inflammation and recovery processes"

### Supplemental Figure 1

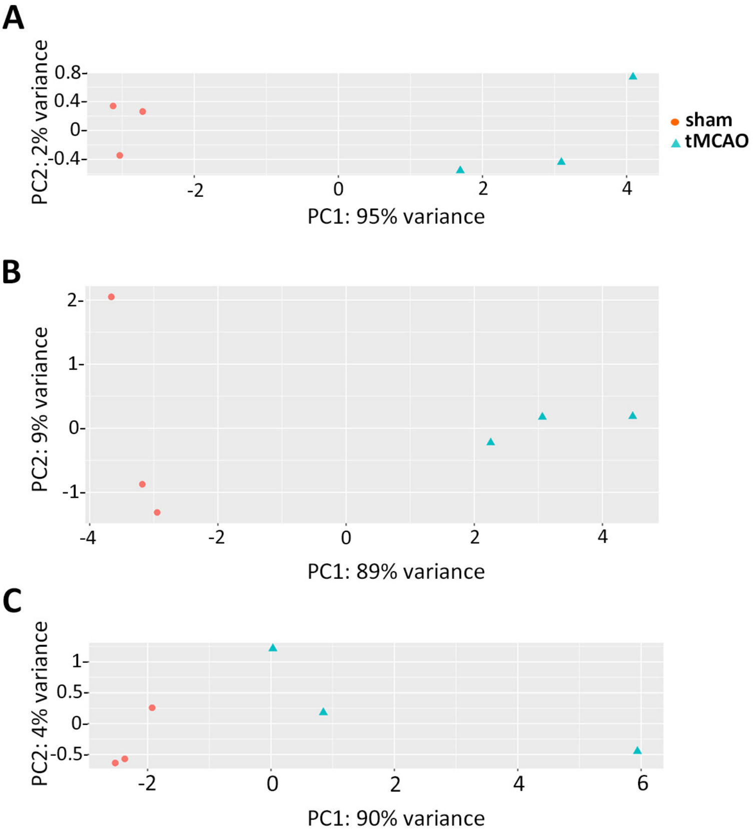

### Supplemental Figure 2

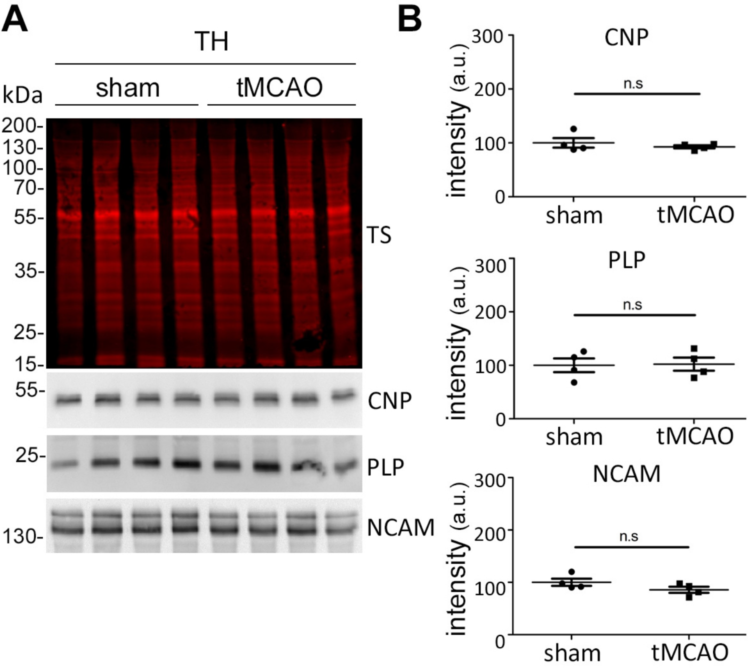

### Supplemental Figure 3

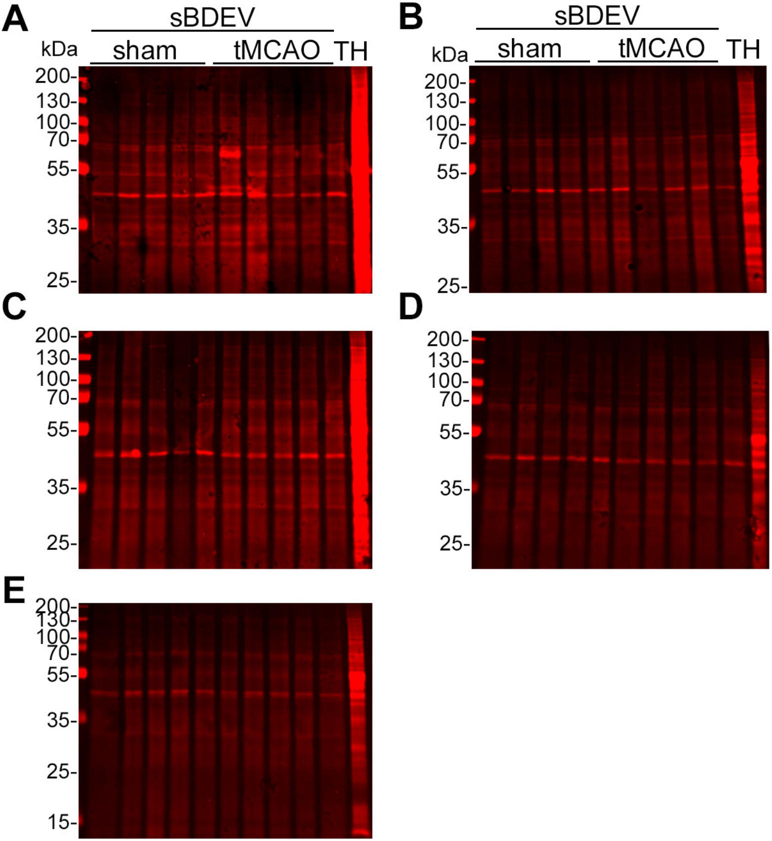
